# Cryo-EM structures of the human G-protein coupled receptor 1 (GPR1) – Gi protein complexes bound to the full-length chemerin and to its C-terminal nonapeptide

**DOI:** 10.1101/2023.06.03.543554

**Authors:** Aijun Liu, Yezhou Liu, Geng Chen, Wenping Lyu, Fang Ye, Junlin Wang, Qiwen Liao, Lizhe Zhu, Yang Du, Richard D. Ye

## Abstract

Chemerin is an adipokine with chemotactic activity to a subset of leukocytes. Chemerin mainly acts through its C-terminal nonapeptide (YFPGQFAFS, C9) that bind to three G protein-coupled receptors including chemokine-like receptor 1 (CMKLR1), G-protein coupled receptor 1 (GPR1) and C-C chemokine receptor-like 2 (CCRL2). We examined C9 signaling through GPR1 and found this receptor capable of Gi signaling but very weak β-arrestin signaling. Here we report high-resolution cryo-EM structures of GPR1-Gi complexes bound to full-length chemerin and to C9, respectively. Unlike C9 that inserts directly into a transmembrane binding pocket, full-length chemerin uses its N-terminal globular core for extensive interaction with the N terminus of GPR1. Within the binding pocket, the C terminal loop containing the nonapeptide takes the same “S-shape” pose as synthetic C9 nonapeptide. These findings explain why the nonapeptide is a full agonist of GPR1, and demonstrate that chemerin uses a “two-site” model for interaction with GPR1. An analysis of the GPR1-Gi protein interface found high similarities to the CMKLR1-Gi complex, as confirmed by site-directed mutagenesis with functional verifications. Our structural analysis demonstrates shared features with chemokines in that chemerin acts as a “reverse chemokine” with switched functions of its N and C termini in the interaction with GPR1.

## Introduction

Chemerin is a small protein encoded by the retinoic acid receptor responder 2 (*RARRES2*) gene. Chemerin is mainly expressed in adipose tissue, liver, lung and skin (Bozaoglu et al., 2007; Goralski et al., 2007; Lehrke et al., 2009). The role of chemerin was initially identified as a chemoattractant of inflammatory cells following its discovery in psoriasis samples (Kennedy & Davenport, 2018). Chemerin was subsequently found to act as an adipokine (Bozaoglu et al., 2007; Goralski et al., 2019; Goralski et al., 2007; Helfer & Wu, 2018). Chemerin secretion and processing requires the removal of the N-terminal signaling peptides of 20 amino acids, resulting in pro-chemerin (amino acids 21-163) with low biological activity. Further proteolytic removal of 6 amino acids from the C-terminus (158-163) leads to chemerin21-157 with full bioactivity (Wittamer et al., 2003). The C-terminal synthetic fragment of human chemerin (149-157, C9) shows comparable biological activity to chemerin21-157 (Meder et al., 2003; Zabel, Allen, et al., 2005; Zabel, Silverio, et al., 2005).

To date, three chemerin receptors have been identified, namely chemokine-like receptor 1 (CMKLR1), G protein-coupled receptor 1 (GPR1) and C-C chemokine receptor-like 2 (CCRL2) (Barnea et al., 2008; Zabel et al., 2008). CMKLR1 responds to chemerin and C9 peptide with activation of the Gi protein pathway and the β-arrestin pathway. In contrast, CCRL2 binds chemerin but does not mediate transmembrane signaling (Kennedy & Davenport, 2018). The biological functions of GPR1 as a chemerin receptor remain unclear. GPR1 was initially identified as an orphan receptor (Marchese et al., 1994). It was subsequently found as a chemerin receptor with uncharacterized pharmacological properties. Using a reporter assay (TANGO) for measurement of β-arrestin activation, Barnea and colleagues found that GPR1 activation is biased towards the β-arrestin pathway when compared with CMKLR1 (Barnea et al., 2008). Another study showed that both CMKLR1 and GPR1 could activate the β-arrestin pathway, but the amplitude of the CMKLR1-mediated signaling was greater (De Henau et al., 2016). In terms of G protein activation, published studies showed downstream activities of RhoA/ROCK, Gα_q/11_ and Gα_i/o_, but it was not clear which one is the dominant G protein for functional coupling (Rourke et al., 2015). Given the inconsistency of these studies and the implications of GPR1 in human immunodeficiency virus replication (Samson et al., 1998; Shimizu et al., 2009; Tokizawa et al., 2000), glucose homeostasis, cardiovascular diseases, steroid hormone synthesis and reproductive biology (Caulfield et al., 2003; Ernst & Sinal, 2010; Karagiannis et al., 2013; Kennedy & Davenport, 2018; Neves et al., 2018; Rourke et al., 2014), it is important to further investigate the structure-function relationship of GPR1.

We recently reported the cryo-EM structure of CMKLR1-Gi complex bound to the C9 peptide of chemerin (Wang et al., 2023), which illustrates a clearly defined binding pocket for the C-terminal peptide of chemerin as well as an interface for Gi protein interaction. GPR1 and CMKLR1 share high sequence homology, but whether the binding event can be translated into G protein activation is unclear. Moreover, the structure of full-length chemerin has not been determined experimentally, nor is its interaction mode with the receptors. To this end, we have solved the cryo-EM structures of GPR1-Gi complex bound to chemerin and its C9 peptide, respectively. Our results revealed the structural basis for GPR1-dependent Gi signaling as well as a chemokine-like “two-site” model of interaction between chemerin and GPR1.

## Methods

### Expression vector design

Human GPR1 was cloned into a pFastBac vector (Invitrogen, Carlsbad, CA) for protein expression and purification. Specifically, the coding sequence of human GPR1 was fused with an N-terminal HA signal peptide followed by a FLAG tag, a human rhinovirus 14 3C (HRV-3C) protease cleavage site (LEVLFQGP) and the thermostabilized apocytochrome b(562)RIL (BRIL) fusion protein (Chun et al., 2012). The coding sequence of human chemerin except for the last six amino acids was synthesized (GENERAL BIOL) and cloned into a pFastbac vector. Human dominant negative Gαi1 (DNGαi1), generated with the G203A and A326S substitution, was cloned into a pFastBac vector. N-terminal 6×His-tagged Gβ1 and Gγ2 were cloned into a pFastBac-Dual vector. scFv16 was fused with an N-terminal GP67 signal peptide and a C-terminal 8× His tag, and the coding sequence was then cloned into a pFastBac vector.

For functional assays, the full-length human GPR1 cDNA were cloned into pcDNA3.1(+) vector (Invitrogen) with an N-terminal FLAG tag. Point mutations were introduced using homologous recombination. Two fragments of GPR1 separated at mutated positions were amplified using PCR and then assembled into pre-cut pcDNA3.1(+) vectors using the ClonExpress Ultra One Step Cloning Kit (Vazyme Biotech; C115). Plasmids with GPR1 mutations were confirmed by DNA sequencing (GENEWIZ).

### Expression and purification of the GPR1-Gi complexes

The baculoviruses expressing GPR1, chemerin, DNGαi1, Gβ1 and Gγ2 were generated and amplified using the Bac-to-Bac baculovirus expression system. The *Sf*9 cells were cultured in SIM SF Expression Medium (Sino Biological). When the cell density reached 3.5 × 10^6^ cells/mL (in total 2 liters), the baculoviruses (GPR1, DNGαi1, Gβ1γ2) were co-expressed in *Sf*9 cells at a ratio of 1:4:2 for the C9-GPR1-Gi complex. the baculoviruses (GPR1, chemerin, DNGαi1, Gβ1γ2) were co-expressed in Sf9 cells at a ratio of 1:2:4:2 for the chemerin-GPR1-Gi complex. After infection for 60 hrs, the cells were collected by centrifugation at 2,000 × g for 15 mins and kept frozen at −80°C before complex purification.

For the purification of C9-bound GPR1-Gi protein complexes, cell pellets from a 2L culture were resuspended in 150 mL lysis buffer (10 mM Tris pH 7.5, 1 mM EDTA, 2.5 μg/mL leupeptin and 160 μg/mL benzamidine, 4 μM C9 peptide and 1 mg/mL iodoacetamide) for 30 mins at room temperature. The lysate was centrifuged for 15 mins at 18,000 × g, and the pellet was homogenized in 150 mL solubilization buffer (20 mM HEPES pH 7.5, 100 mM NaCl, 10% glycerol, 1% dodecylmaltoside (DDM), 0.1% cholesteryl hemisuccinate (CHS), 2.5 μg/mL leupeptin and 160 μg/mL benzamidine, 4 μM C9 peptide, 1 mg/mL iodoacetamde, 2 mg scFv16, 25 mU/mL apyrase) using a Dounce homogenizer. The sample was stirred for 2 hrs at 4°C and then centrifuged for 30 mins at 18,000 × g to remove the insoluble debris. The solubilized supernatant fraction was incubated with 2 mL anti-FLAG affinity resin (GenScript Biotech, Piscataway, NJ) and stirred at 4°C for 2 hrs. Then, the resin was manually loaded onto a gravity-flow column and extensively washed with the FLAG wash buffer (W1: 20 mM HEPES pH 7.5, 0.1% DDM, 0.01% CHS, 100 mM NaCl, 2 mM CaCl_2_, 4 μM C9 peptide. W2: 20 mM HEPES pH 7.5, 0.2% lauryl maltose neopentyl glycol (LMNG), 0.02% CHS, 100 mM NaCl, 2 mM CaCl_2_, 4 μM C9 peptide) by mixing W1 and W2 buffer in the following ratios: 5 mL:5 mL, 2 mL:8 mL, 1 mL:9 mL, 0.5 mL:9.5 mL, 0 mL:10 mL, respectively. The GPR1-Gi complexes attached to the resin were further eluted with 10 mL elution buffer (20 mM HEPES pH 7.5, 0.01% LMNG, 0.002% CHS, 100 mM NaCl, 4 μM C9 peptide, 5 mM EDTA, 0.2 mg/ml FLAG peptide). Eluted protein complexes were concentrated to 400 μL in an Amicon® Ultra-15 Centrifugal Filter Unit (Millipore, Burlington, MA) and further subjected to a size exclusion chromatography through a Superdex 200 Increase 10/300 column (GE Healthcare Life Sciences, Sweden) equipped in an AKTA FPLC system with running buffer (20 mM HEPES pH 7.5, 0.01% LMNG, 0.002% CHS, 100 mM NaCl, 4 μM C9 peptide). Eluted fractions containing GPR1-Gi complexes were re-pooled and concentrated before being flash frozen in liquid nitrogen and stored at −80°C. The purification process of the chemerin-GPR1-Gi complex was almost identical except C9 was not added.

### Expression and purification of scFv16

The antibody fragment scFv16 was expressed as a secretory protein and purified as previously described (Chen et al., 2022). Briefly, *Trichoplusia ni* Hi5 insect cells were cultured to reach a density of 3.5 × 10^6^ cells/mL. Cells were then infected with scFv16 baculovirus at a ratio of 1:50. After 60 hrs of culture, the supernatant was collected and loaded onto a Ni-NTA resin column. The column was washed with 20 mM HEPES (pH 7.5), 500 mM NaCl, and 20 mM imidazole, and then subjected to elution by 20 mM HEPES (pH 7.5), 100 mM NaCl, and 250 mM imidazole. The eluted proteins were concentrated and subjected to size-exclusion chromatography using a Superdex 200 Increase 10/300 column (GE Healthcare). Finally, the purified scFv16 protein with a monomeric peak was concentrated and flash frozen in liquid nitrogen and stored at −80 °C for further use.

### Cryo-EM sample preparation and data collection

For cryo-EM sample preparation of the C9-GPR1-Gi-scFv16 complex or chemerin-GPR1-Gi complex, UltrAuFoil Au R1.2/1.3 300-mesh grids (Quantifoil) was glow discharged in a Tergeo-EM plasma cleaner. 3 μL purified complex sample was loaded on the grid and blotted for 3 s with a blotting force of 0, and then flash-frozen in liquid ethane cooled by liquid nitrogen using Vitrobot Mark IV (Thermo Fisher Scientific, Waltham, MA). Cryo-EM data were collected at the Kobilka Cryo-EM Center of The Chinese University of Hong Kong, Shenzhen, on a 300 kV Titan Krios Gi3 microscope (Thermo Fisher Scientific). The raw movies were recorded using a Gatan K3 BioQuantum Camera at the magnification of 105,000, with a pixel size of 0.83 Å. A GIF Quantum energy filter was applied to exclude inelastically scattered electrons (Gatan) using a slit width of 20 eV. The movie stacks were acquired with a total exposure time of 2.5 s fragmented into 50 frames (0.05 s/frame). The defocus range was from −1.2 to −2.0 μm. The semi-automatic data acquisition was performed using SerialEM. A total of 3,609 image stacks were collected in 48 hrs for the C9-GPR1-Gi-scFv16 complex, and a total of 7,706 image stacks were collected in 72 hrs for the chemerin-GPR1-Gi-scFv16 complex.

### Image processing and model building

Data processing was performed with cryoSPARC 3.3.1 (Structura Biotechnology Inc., Toronto, Canada). Patch motion correction and patch CTF estimation were applied to the image stacks. For the chemerin-GPR1-Gi-scFv16 dataset, 2,232,191 particles were auto-picked and subjected to 2D classification. Ab initio reconstruction was performed by selecting particles in good 2D classes. The high-quality particles were selected by multiple rounds of heterogeneous refinements. Finally, a dataset with 107,273 particles was used for non-uniform refinement and local refinement, yielding a final map with a global resolution at 3.29 Å by FSC 0.143 cutoff criterion. For the C9-GPR1-Gi-scFv16 dataset, 2,280,697 particles were auto-picked. 2D classification was performed, resulting in 732,629 particles selected for *ab initio* reconstruction. After 3 rounds of heterogeneous refinements, a final set of 339,859 particles was exported to non-uniform refinement and local refinement, yielding a map with a global resolution of 2.90 Å.

The predicted structure GPR1 and chemerin on the AlphaFold database was used to build the initial model. The coordinates of Gi1 and scFv16 from the CMKLR1-Gi-Scfv16 complex (PDB ID: 7YKD) were applied as templates. All models were docked into the EM density map using UCSF Chimera version 1.12, followed by iterative manual building in Coot-0.9.2 and refinement in Phenix-1.18.2. The statistics of the final model were further validated by Phenix-1.18.2. Structure figures were generated by Chimera or PyMOL (Schrödinger, Inc., New York, NY). The statistics of data-collection and structure-refinement are shown in Table S1.

### Molecular modeling and molecular dynamic simulation

The protonation state of the GPR1 was assigned by the web server H++ (Anandakrishnan et al., 2012) assuming pH 7.4, and CHARMM36m (Anandakrishnan et al., 2012) force field was employed in all simulations. After energy minimization, membrane relaxation, and equilibrium simulation (Huang et al., 2017), ten independent 1-µs long production MD simulations were carried out for C9-GPR1 binding. Fifty-thousand conformations were collected in total from the assemble of trajectories. Hydrogen bonds were identified based on cutoffs for the Donor-H···Acceptor distance and angle. The criterion employed was angle > 120° and H···Acceptor distance < 2.5 Å in at least 10% of the trajectory.

### G protein dissociation assay

G protein activation was tested by a NanoBiT-based G protein dissociation assay (Inoue et al., 2019). HEK293T cells were plated in a 24-well plate 24 hrs before transfection. Lipofectamine™ 3000 (Invitrogen, L3000001) transfection was performed with a mixture of 92 ng pcDNA3.1 vector encoding human GPR1 (WT/mutants) or WT human CMKLR1 for comparison, 46 ng pcDNA3.1 vector encoding Gαi1-LgBiT, 230 ng pcDNA3.1 vector encoding Gβ1 and 230 ng pcDNA3.1 vector encoding SmBiT-Gγ2 (per well in a 24-well plate), respectively. After 24 hrs of incubation, the transfected cells were collected and resuspended in HBSS containing 20 mM HEPES. The cell suspension was loaded onto a 384-well culture white plate (PerkinElmer Life Sciences, Waltham, MA) at a volume of 20 μL and loaded with 5 μL of 50 μM coelenterazine H (Yeasen Biotech, Shanghai, China). After 2 hrs of incubation at room temperature, the baseline was measured using an Envision 2105 multimode plate reader (PerkinElmer). Then, full-length chemerin (abcam, Cambridge, UK; ab256228) or synthesized C9 peptides (ChinaPeptides, Shanghai, China) were added to the cells in different concentration. The ligand-induced luminescence signals were measured 15 mins after ligand addition and divided by the initial baseline readouts. The fold changes of signals were further normalized to PBS-treated signal and the values (EC_50_) were expressed as a function of different ligand concentrations based on three independent experiments, each with triplicate measurements.

### cAMP assay

Wild-type human GPR1 and its mutants, or wild-type human CMKLR1 for comparison, were transiently expressed in HeLa cells 24 hrs prior to collection. The cells were resuspended in HBSS buffer plus 5 mM HEPES, 0.1% BSA (w/v) and 0.5 mM 3-isobutyl-1-methylxanthine and loaded onto 384-well plates. Different concentrations of full-length chemerin or C9 peptide were prepared with 2.5 μM forskolin in the abovementioned buffer. The cells were stimulated by the ligands and 2.5 μM forskolin for 30 mins in a cell incubator. Intracellular cAMP levels were measured with the LANCE Ultra cAMP kit (PerkinElmer, TRF0263) following the manufacturer’s instructions. In the measurements, signals of time resolved-fluorescence resonance energy transfer (TR-FRET) were detected by an EnVision 2105 multimode plate reader (PerkinElmer). Intracellular cAMP levels were calculated according to the TR-FRET signals of the samples and cAMP standards.

### β-arrestin recruitment assay

For NanoBiT-based β-arrestin recruitment assays, HEK293T cells were seeded in a 24-well plate 24 hrs before transfection. Cells are co-transfected with GPR1-WT-smBiT or CMKLR1-WT-smBiT (400 ng/well) and LgBiT-β-Arr1 or LgBiT-β-Arr2 (200 ng/well) by Lipofectamine™ 3000 (Invitrogen) for 24 hrs. Cells were collected and resuspended in HBSS buffer containing 20 mM HEPES, and then 20 μL of cell suspension was loaded onto a 384-well white plate at a concentration of 2×10^4^ cells/well. Test samples were further loaded with coelenterazine H to a final concentration of 10 μM. After 25 mins incubation at 37 °C, the samples were measured for baseline luminescence using an Envision 2105 multimode plate reader (PerkinElmer). Different concentrations of full-length chemerin or C9 peptide were added to the wells and the luminescence signals were detected for 30 mins. The signal readouts were further normalized to PBS-treated signal and the values (EC_50_) were expressed as a function of different ligand concentrations based on three independent experiments, each with triplicate measurements.

### IP1 accumulation assay

Wild-type GPR1 and its mutants, and wild-type human CMKLR1 for comparison, were transiently expressed in HEK293T cells for 24 hrs. IP1 accumulation was tested using the IP-One Gq HTRF kit (Cisbio). The cells were resuspended in the stimulation buffer (Cisbio) and incubated with different concentrations of C9 peptide diluted in the stimulation buffer for 30 mins at 37°C. The accumulation of IP1 was further determined following the manufacturer’s protocols. Fluorescence intensities were measured on an Envision 2105 multimode plate reader (PerkinElmer). Intracellular IP1 levels were calculated according to the fluorescence signals of the samples and IP1 standards.

### GPR1 expression level determination by flow cytometry

HEK293T cells were transfected with FLAG-tagged WT or mutant GPR1 expression plasmids for 24 hrs at 37°C. Then the cells were harvested and washed in HBSS containing 5% BSA for three times on ice. The cells were then incubated with a FITC-labeled anti-FLAG antibody (Sigma, Cat #F4049; 1:50 diluted by HBSS buffer) for 30 mins on ice and washed with HBSS. The FITC fluorescence signals demonstrating the antibody-receptor complex on the cell surface were quantified by flow cytometry (CytoFLEX, Beckman Coulter). Relative expression levels of GPR1 mutants were represented according to the fluorescence signals.

### Statistical analysis

The data were analyzed with Prism 9.5.0 (GraphPad, San Diego, CA). For dose-response analysis, the curves were plotted with the log[agonist] vs. response equation (three parameters) in the software. For cAMP, IP1, and G protein dissociation assays, data points were presented as the percentages (mean ± SEM) of the maximal response level for each sample, from at least three independent experiments, as indicated in figure legends. For the β-arrestin recruitment assay, data were presented as raw chemiluminescence signals (mean ± SEM) from at least three independent experiments. The EC_50_ values were obtained from the dose-response curves. For cell surface expression, data points were presented as the percentages (mean ± SEM) of the flow cytometry fluorescence signals of WT GPR1. For statistical comparisons, Analysis of Variance (ANOVA) was performed using the one-way method. A *p*-value of 0.05 or lower is considered statistically significant.

## Results

### GPR1 functionally couples to the Gi proteins

It has been long controversial about the downstream signaling events elicited by GPR1. We hence compared transmembrane signaling through GPR1 and CMKLR1 by performing G protein dissociation assay, cAMP inhibition assay measuring Gαi response, IP1 accumulation assay for Gβγ signaling, and β-arrestin recruitment assays in GPR1-expressing cells stimulated with the C9 peptide at different concentrations. GPR1 elicited cAMP inhibition, although to a lesser extent than CMKLR1 (Fig. 1A). In NanoBiT-based G protein dissociation assay, C9 activated GPR1 and CMKLR1 with a similar EC_50_, although the efficacy was lower for GPR1 (Fig. 1B). In IP1 accumulation assay, a higher EC_50_ with a similar amplitude was observed for GPR1 (Fig. 1C). These results demonstrate Gi coupling activities of GPR1 in response to C9 stimulation. We next examined β-arrestin recruitment. Both CMKLR1 and GPR1 preferentially recruited β-arrestin1 over β-arrestin2 (Fig. 1D). However, GPR1 experienced a significantly lower efficacy for β-arrestin1 recruitment, with nearly no recruitment of β-arrestin2. These findings demonstrate that the major signaling differences between GPR1 and CMKLR1 lie in the recruitment of β-arrestins rather than G protein activation in contrast to previous reports (Barnea et al., 2008; De Henau et al., 2016; Degroot et al., 2022; Fischer et al., 2021; Rourke et al., 2015; Wittamer et al., 2003).

**Figure 1.**
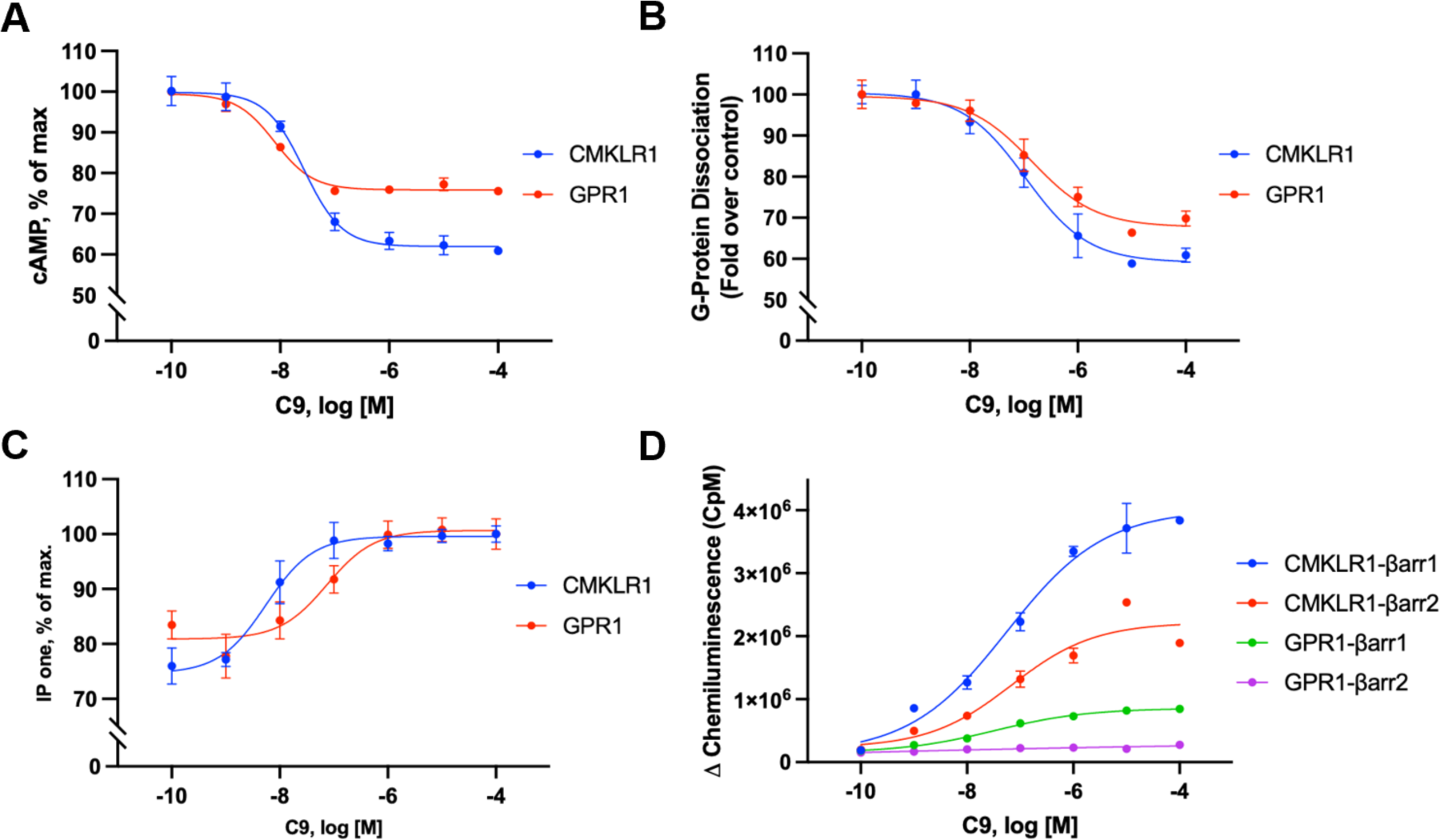
Gi signaling mediated by GPR1 and CMKLR1. **(A)** cAMP inhibition mediated by GPR1 and CMKLR1 in response to different concentrations of the C9 peptide. (B) G protein dissociation assay based on NanoBiT technology in transfected cells that express GPR1 and CMKLR1, respectively, and stimulated with C9 at different concentrations. **(C)** IP-one accumulation in cells expressing GPR1 and CMKLR1, respectively, in response to C9 at different concentrations. (D) |3-arrestin recruitment upon 09 stimulation based on NanoBiT technology, in transfected cells expressing GPR1 and CMKLR1, respectively. Data shown are means ± SEM from three independent experiments.

### Cryo-EM structure of the C9-GPR1-Gi complex

Having confirmed GPR1 coupling to Gi in transfected cells, we sought to determine physical interactions between GPR1 and G proteins using purified receptor and Gi proteins. As shown in Fig. S1, complex formation was readily observed between the C9-bound GPR1, the Gi heterotrimer and the scFv16 antibody fragment that was used for stabilization of the Gi-containing complex (Maeda et al., 2018). The cryo-EM structure of the C9-bound GPR1-Gi complex was next determined to an overall resolution of 2.90 Å (Fig. 2A, 2B, Fig. S1).

**Figure 2.**
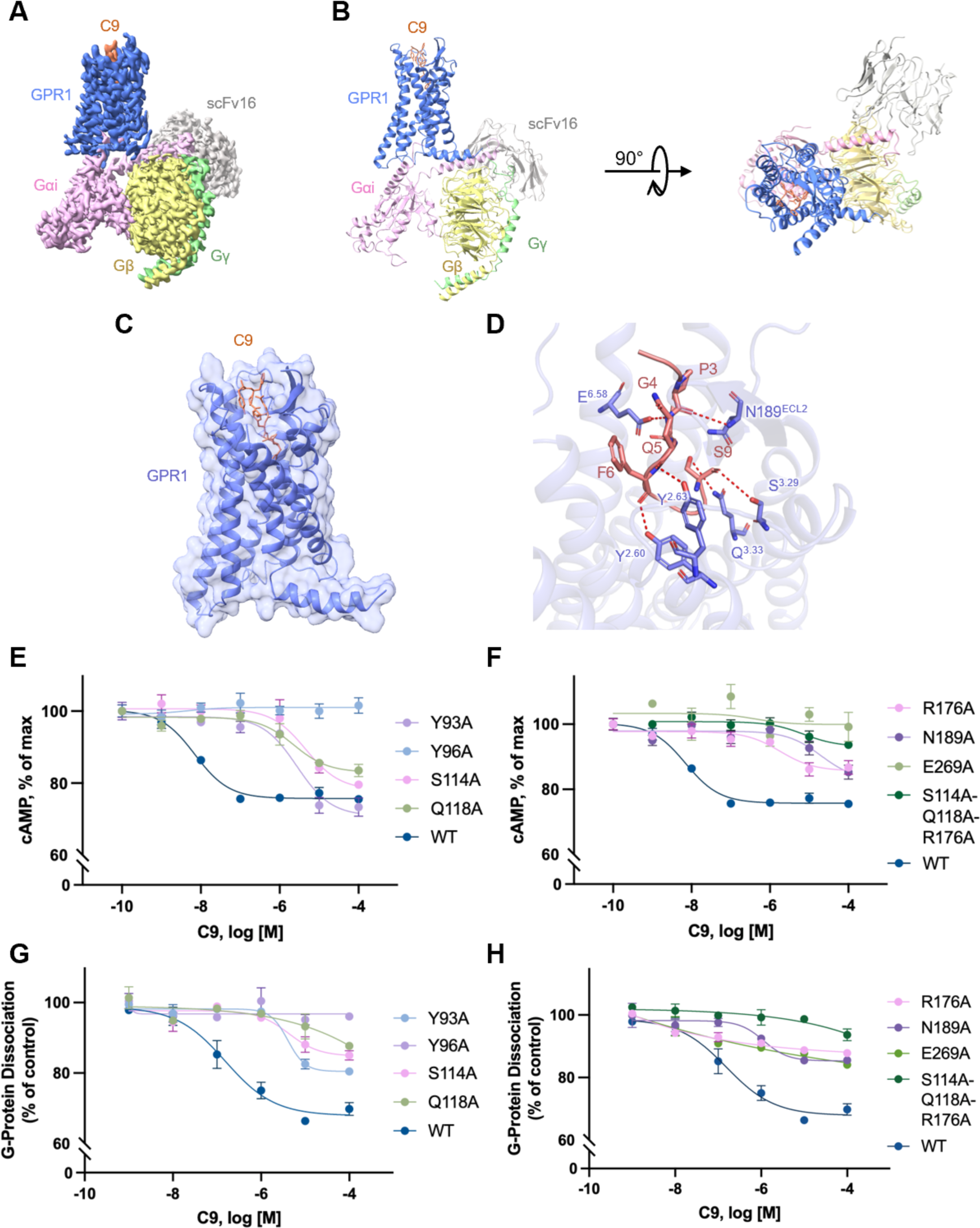
The structural model of C9-GPR1-Gi complex. **(A)** Cryo-EM density map of the C9-bound GPR1-Gi-scFv16 complex. **(B)** Overall structure of C9-GPR1-Gi-scFv16 complex from side view (left) and extracellular view (right). **(C)** Interaction between C9 and GPR1. The receptor is shown by cartoon and surface representation. 09 is shown in salmon color. (D) Polar interactions between C9 and GPR1. The hydrogen bonds are displayed as dashed lines. The residue numbering of GPR1 follows the Ballesteros-Weinstein nomenclature. **(E-F)** Effects of alanine substitution of selected amino acids in the C9 binding pocket on cAMP inhibition in cells expressing the GPR1 mutants. **(G-H)** The effects on G protein dissociation of the same set of GPR1 mutants in transfected cells stimulated with different concentrations of C9. Data shown are means + SEM from three independent experiments.

The C9 peptide fit snugly in a transmembrane (TM) pocket formed by TM2, TM3, TM4, TM6 and TM7. The N-terminal end of the peptide was barely visible when the complex was viewed from the side. Within the binding pocket, the C9 peptide assumed an “S”-shape (Fig. 2 C and D). The N-terminal Y1 (corresponds to Y149 in the full-length chemerin) and F2 of the C9 peptide showed extensive hydrophobic interactions with L186^ECL2^, H273^ECL3^, Y188^ECL2^ and I272^6.61^. P3 of the C9 peptide formed polar interaction with N189^ECL2^. G4 had polar interaction formed between its backbone amide group and E269^6.58^. Q5 with its backbone carbonyl oxygen had a polar interaction with Y96^2.63^. The aromatic ring of F6 showed nonpolar interactions with F101^ECL1^ and Q283^7.32^, and its backbone carbonyl oxygen had polar interaction with Y93^2.60^. F8 with its aromatic ring showed extensive hydrophobic interactions with residues P287^7.36^, I286^7.35^, C187^ECL2^, T290^7.39^, A117^3.32^ and M121^3.36^ to stabilize the peptide ligand in the binding pocket. S9 at the C-terminal end of the C9 peptide experienced polar interactions extensively with S114^3.29^, Q118^3.33^ and R176^4.64^, through oxygens in its carbonyl and side chain. Of note, Q^3.33^ is conserved in some chemoattractant GPCRs including FPR1 and FPR2 (Chen et al., 2022; Chen et al., 2020; Zhu et al., 2022; Zhuang et al., 2020; Zhuang et al., 2022), suggesting that GPR1 shares structural features in ligand binding with chemotactic peptide receptors.

Alanine substitutions of GPR1 residues forming polar interactions with the C9 peptide including Y93^2.60^, Y96^2.63^, S114^3.29^, Q118^3.33^, R176^4.64^, N189^ECL2^, and E269^6.58^, resulting in a remarkable decrease in the potency of the C9 peptide (Fig. 2 E-H). Among these substituted residues, Y96^2.63^A and E269^6.58^A completely diminished signaling triggered by the C9 peptide. Due to extensive polar interactions between S9 in the C9 peptide and several amino acids in GPR1, single point mutations of S114^3.29^, Q118^3.33^ or R176^4.64^ did not fully eliminate the response. This result can be explained by the flexibility at S9 where multiple hydrogen bonds may form dynamically between the carbonyl and side chain oxygens of S9 and S114^3.29^, Q118^3.33^ and R176^4.64^ of GPR1. Our prediction was confirmed after introducing triple alanine substitutions at S114/Q118/R176, that completely abolished the agonist-induced cAMP inhibition and G protein dissociation upon C9 stimulation.

### Cryo-EM structure of the full-length chemerin-GPR1-Gi complex

We next examined the full-length chemerin for its structure and its mode of interaction with GPR1. It is especially intriguing why, of the 137 a.a. mature chemerin, only 9 would be sufficient for full agonistic activity. To this end, we determined the cryo-EM structure of the full-length chemerin-GPR1-Gi complex to an overall resolution of 3.29 Å (Fig. 3 A and B, Fig. S2 and S3). Except the N terminal 13 amino acids, GPR1 in this complex is clearly defined. The majority of the full-length chemerin sits on top of the TM pocket, with interaction between the N terminal β-strand of GPR1 and the globular core of chemerin (Fig. 3B, 3C). This mode of interaction differs from most of chemokine receptors that have their N terminal fragments wrapping around the globular core of chemokines (Kufareva et al., 2017; Urvas & Kellenberger, 2023). Analysis of the chemerin structure, that was not experimentally determined before, identified an N-terminal α helix (H1) followed by 4 anti-parallel β-strands (β1-β4) and another α helix (H2) in parallel with the ý-strands (Fig. 3D). The extruding C-terminal tail forms a C9 loop that extends deep into the TM binding pocket (red in Fig. 3D). The “S-shape” of the C9 loop is clearly visible in the binding pocket similar to the C9 peptide (Fig. 3E).

**Figure 3.**
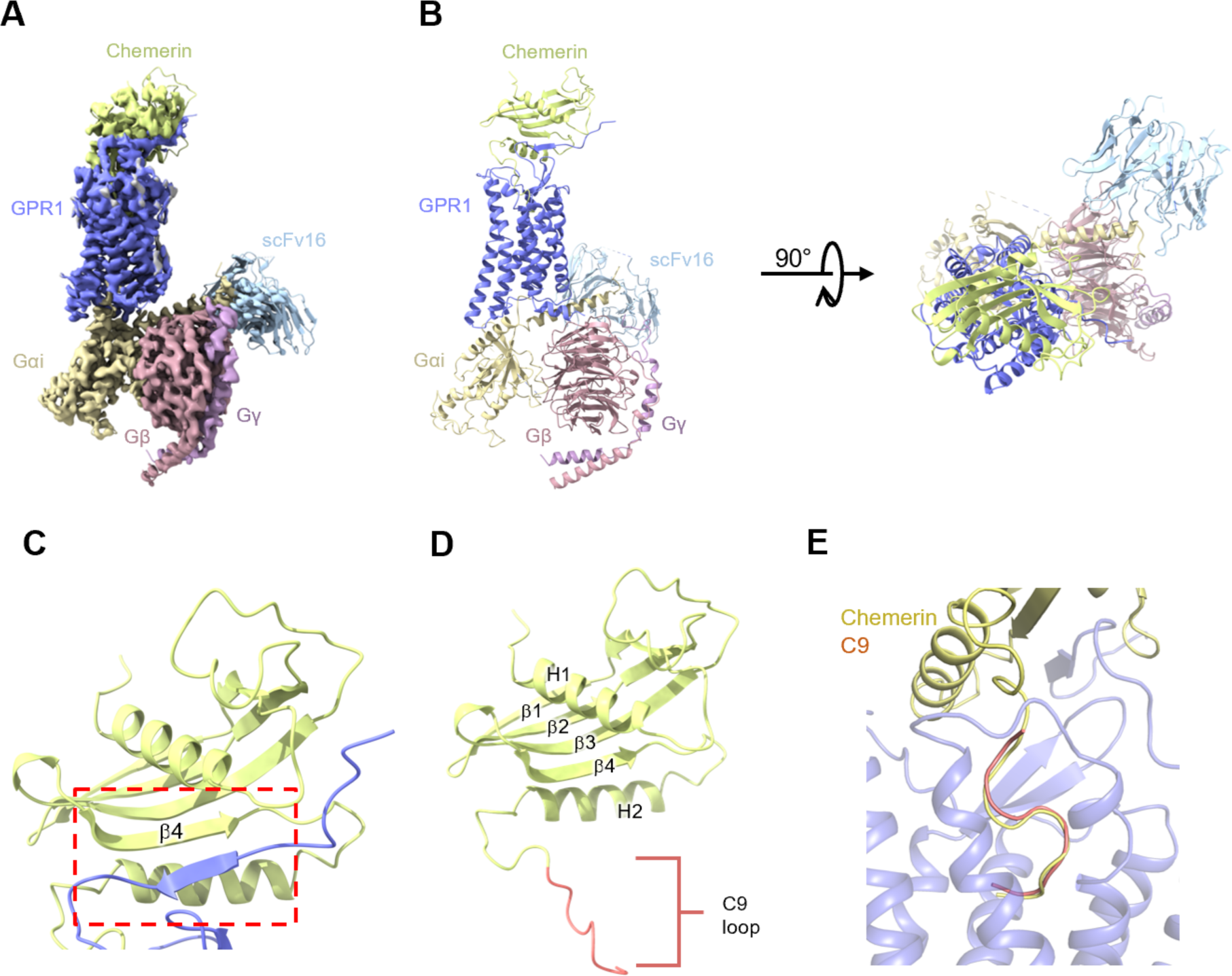
Overall structure of full-length chemerin-GPR1-Gi complex. **(A)** Cryo-EM density map of GPR1-Gi-scFv16 complex bound to chemerin. **(B)** Overall structure of full-length chemerin-GPR1-Gi-scFv16 complex from side view (left) and extracellular view (right). (C) Interaction between the p4 strand of chemerin (grass green) and the N-terminal p strand of GPR1 (slate blue). (D) Structure of full-length chemerin (21-157). Structural motifs including helices H1 and H2, p-strands p1 −4 and the C-terminal C9 loop (red). (E) Comparison between the binding poses of chemerin and C9 peptide in the TM pocket of GPR1. Chemerin is shown in yellow, and C9 peptide is shown in salmon.

In the structure of the chemerin-bound GPR1, multiple binding regions are present between the chemerin ligand and the receptor. There is chemerin binding region 1 (CBR1) at the N-terminus of the receptor that interacts with the β4 strand of chemerin (Fig. 4A). There is also a chemerin binding region 2 (CBR2), defined by the TM binding pocket that interacts only with the C9 loop of chemerin (Fig. 4A). Detailed molecular interactions between the full-length chemerin and GPR1 were analyzed (Fig. 4B). Multiple polar interactions were observed at CBR1, where S23^NT^ and E25^NT^ interact with R113 of chemerin, and S26^NT^ forms a hydrogen bond with L111 at the β4 strand of chemerin [superscripts indicate the Ballesteros-Weinstein numbering scheme for GPCRs (Ballesteros & Weinstein, 1995)] (Fig. 4B). Multiple non-polar interactions between the proximal N-terminus of GPR1 and the loop between H1 and β1 of chemerin strengthen and stabilize chemerin binding at CBR1. Like C9 interaction with the ligand binding pocket, the canonical TM binding pocket for full-length chemerin is surrounded by TM2, 3, 4, 6, and 7. These structures together form the CBR2. Multiple polar bonds are present between the chemerin C-terminal C9 loop region (126-137) and receptor CBR2 (Fig. 4B). The sidechain of chemerin H146 forms polar bond with I272^6.61^, and Y149 interacts with E269^6.58^. Similar to C9 peptide binding to GPR1, the backbone carbonyl oxygen of P151 forms hydrogen bond with N189^ECL2^, and polar interactions are present between Q153 and Y96^2.63^. Also in CBR2, F154 uses its backbone carbonyl oxygen to form polar interaction with Y93^2.60^, and the C-terminal S157 has hydrogen bond with Q118^3.33^. F150 with its aromatic ring forms nonpolar interactions with V179^ECL2^, the aromatic ring of F154 has hydrophobic interactions with F101^ECL1^ and Q283^7.32^, and extensive hydrophobic interactions are present between chemerin F156 and GPR1 A117^3.32^, T290^7.39^, S114^3.29^ and R176^4.64^. Of interest, another pair of polar interactions is found between E129 of chemerin at helix H2 and H194^5.26^ of GPR1 near ECL2. A similar polar interaction feature is present between some chemokines and ECL2 of the respective chemokine receptors and defined as chemokine recognition site 3 (CRS3) (Urvas & Kellenberger, 2023), this region of GPR1 can hence be categorized into chemerin binding region 3 (CBR3). These three chemerin binding regions work together to stabilize receptor-chemerin interactions and activate the receptor.

**Figure 4.**
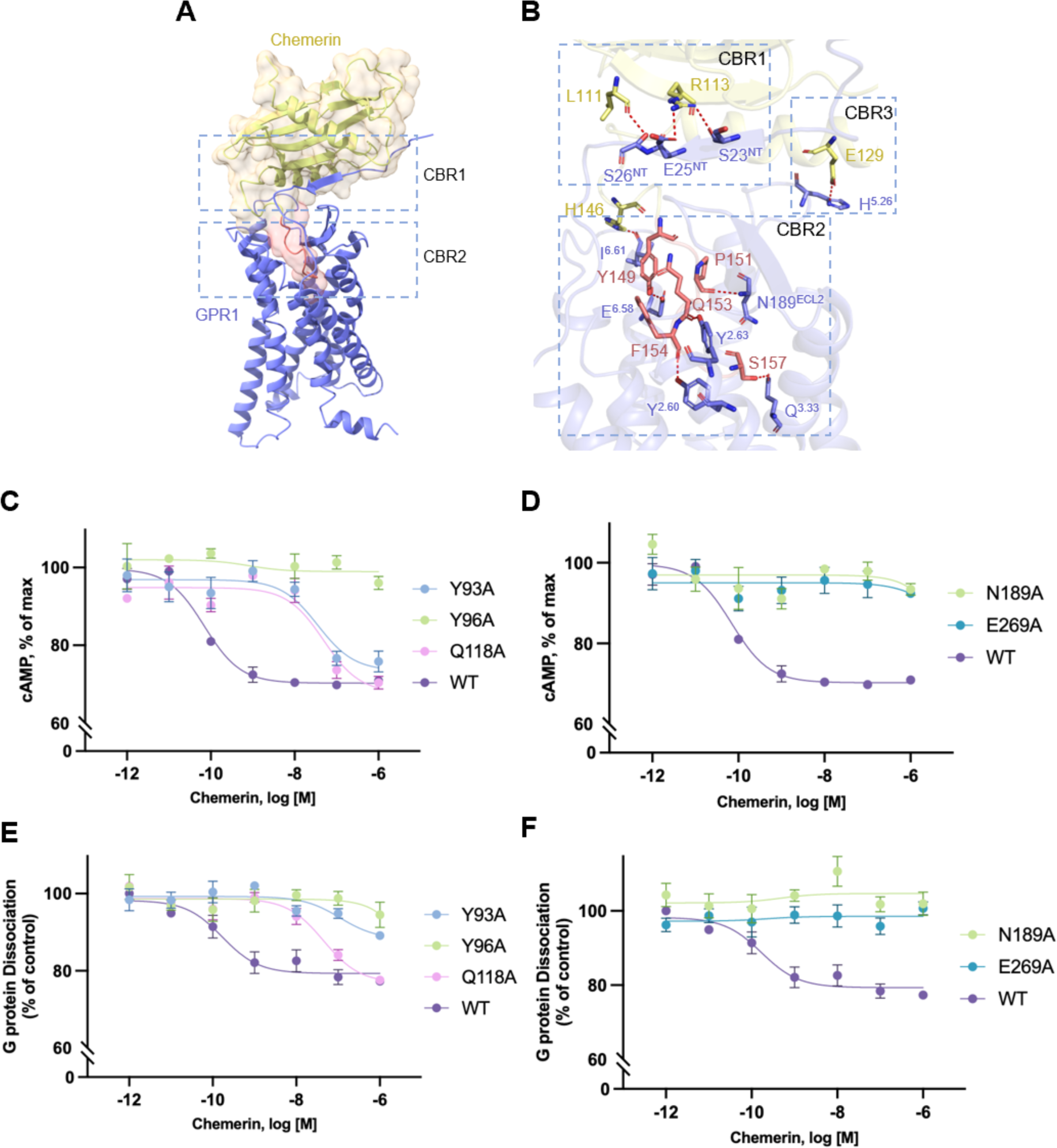
Ligand binding pockets in chemerin-GPR1-Gi complex. **(A)** Interaction between chemerin and GPR1. Chemerin is shown by cartoon and transparent surface representation. The C-terminal nonapeptide (C9) is highlighted in salmon color. **(B)** Polar interactions between chemerin and GPR1. The hydrogen bonds are displayed as dashed lines. The residue numbering of GPR1 follows the Ballesteros-Weinstein nomenclature. (C-D) Effects of alanine substitution of selected amino acids in the chemerin binding pocket on cAMP inhibition, in cells expressing the GPR1 mutants. (E-F) The effects on G protein dissociation of the same set of GPR1 mutants in transfected cells stimulated with different concentrations of chemerin. Data shown are means ± SEM from three independent experiments.

Alanine substitutions of the key residues in the CBR2 binding pocket of GPR1 were performed, and the resulting receptors were expressed for functional assays including cAMP inhibition and G protein dissociation to determine any changes in GPR1 activation by the full-length chemerin (Fig. 4 C-F). The EC_50_ of chemerin in eliciting cAMP inhibition and G protein dissociation was around 10^-10^, while for C9 peptide, the EC_50_ is 10^-8^. In response to chemerin binding, the Y93^2.60^A and Q118^3.33^A mutants reduced the potency by three magnitudes, while Y96^2.63^A, N189^ECL2^A and E269^6.58^A completely abolished both cAMP inhibition and G protein dissociation. All mutants were properly expressed on the cell surface (Fig. S4), thus excluding global misfolding of the receptors after alanine substitution at these positions. Functional assays revealed that, despite having indistinguishable postures in the CBR2 pocket, C9 and the full-length chemerin have different sidechain orientations, leading to slightly altered polar contacts with the receptor TM pocket. Altogether, these results support the role of the substituted amino acid residues of GPR1 in polar interactions with the full-length chemerin.

### Thermodynamic stability of GPR1-C9 interface

Full-atom molecular dynamics (MD) simulations were performed for the GPR1-C9 complex at room temperature (10 replicas of 1-µs-long simulation). In these MD trajectories, the complex captured by cryo-EM were overall stable under thermodynamic perturbation (Fig. S5). The binding pose of the C9 peptide were well kept through the assemble of 1-µs trajectories (Fig. 5 and Fig. S6). Except the hydrophobic Phe (F2), all of the 9 residues in C9 peptide form hydrogen-bonds (H-bonds) with GPR1 (Fig. 5). The residues close to the C-terminal end of the C9 peptide form several H-bonds with Y96^2.63^, N189^ECL2^, R176^4.64^, S114^3.29^, Y262^6.51^, and K210^5.42^. N189^ECL2^ and Y96^2.63^ also form H-bonds with P3 and Q5 on the C9 peptide. The N-terminus of the peptide ligand likely interacts with negatively charged E269^6.58^. The latter also interacts with G4, which forms a stable H-bond with R176^4.64^ (Fig. 5B). These interactions result in a complex network between C9 and the receptor, which rationalizes the thermodynamic stability of the C9 peptide in the binding pocket of GPR1.

**Figure 5.**
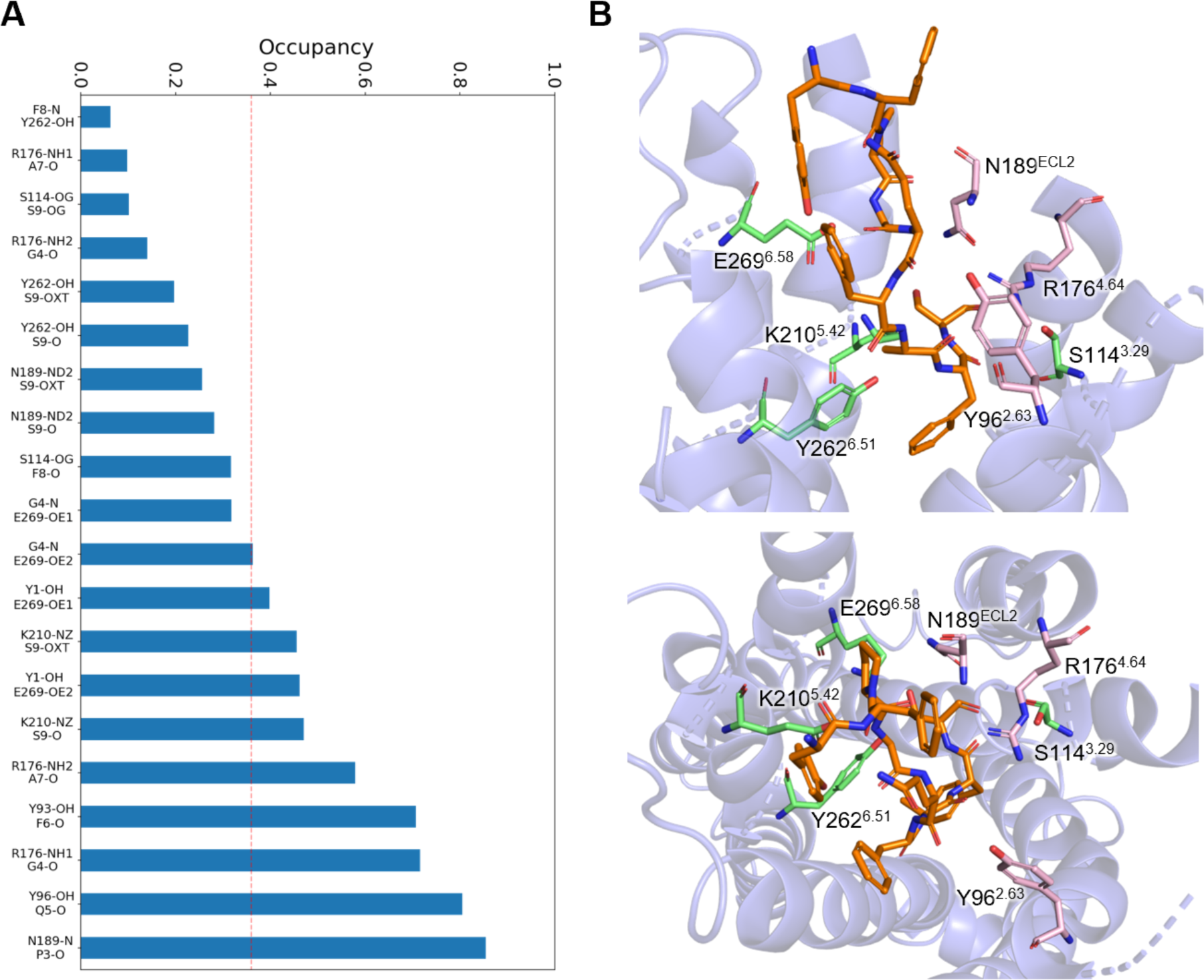
Thermodynamic stability analysis of the GPR1-C9 interface with ps-scale MD simulations. **(A)** The occupancy of all H-bonds between C9 and GPR1 observed in MD simulations. **(B)** Side view (upper panel) and top view (lower panel) of the distributions of functionally related residues around the C9 peptide.

Results of the MD simulations corroborate functional assay data based on the EC_50_ values of cAMP inhibition assays, using WT and alanine substituted mutants of GPR1. Alanine substitution of N189^ECL2^ and Y96^2.63^, which have the top 2 occupancy scores on single H-bond (Fig. 5A), abrogated cAMP inhibition (-logEC_50_ = 0), indicating their functional importance. Another functionally important residue is E269^6.58^, which has low single H-bond occupancy (<0.6, Fig. 5A) but the highest overall H-bond occupancy (occupancy values >1.5) when all associated H-bonds were considered. Its substitution with alanine led to a loss of cAMP inhibition. Therefore, there is a high-level correlation between the polar interactions of C9 with GPR1 and its functions.

### Activation mechanism of the GPR1-Gi complex

To investigate the conformational changes associated with the activation of GPR1, we compared the structure of active GPR1 and an antagonist-bound inactive C5aR (C5aR-PMX53, PDB ID: 6C1R) for its homology to GPR1, or a C9-bound active CMKLR1 (CMKLR1-C9, PDB ID: 7YKD), respectively (Fig. 6). The comparison also revealed an outward movement of TM5 and TM6, and an inward movement of TM7, in the activation of GPR1 (Fig. 6A). Of note, the D^3.49^-R^3.50^-Y^3.51^ motif found in most Class A GPCRs for G protein activation, was replaced with D134-H135-Y136 in GPR1. Previous reports indicate that polar interaction of R^3.50^ and Y^5.58^ is important for G protein activation (Wang et al., 2023; Weis & Kobilka, 2018; Zhou et al., 2019) (Fig. 6B). However, the apparent activation of G proteins in GPR1, which only has H^3,50^ in this motif, suggests that the R^3.50^-Y^5.58^ interaction is not critical to G protein activation. Supporting this notion, our substitution of H^3.50^ with arginine did not significantly change G protein dissociation (Fig. 8D) but made the receptor less capable of inhibiting cAMP (Fig. 8A). Alanine substitution of H^3,50^ led to a complete loss of function (Fig. 8 A and D). These results indicate that the D^3.49^-H^3.50^-Y^3.51^ motif in the context of the overall sequence of GPR1 remains functional in G protein activation, even in the absence of an R^3.50^-Y^5.58^ interaction.

**Figure 6.**
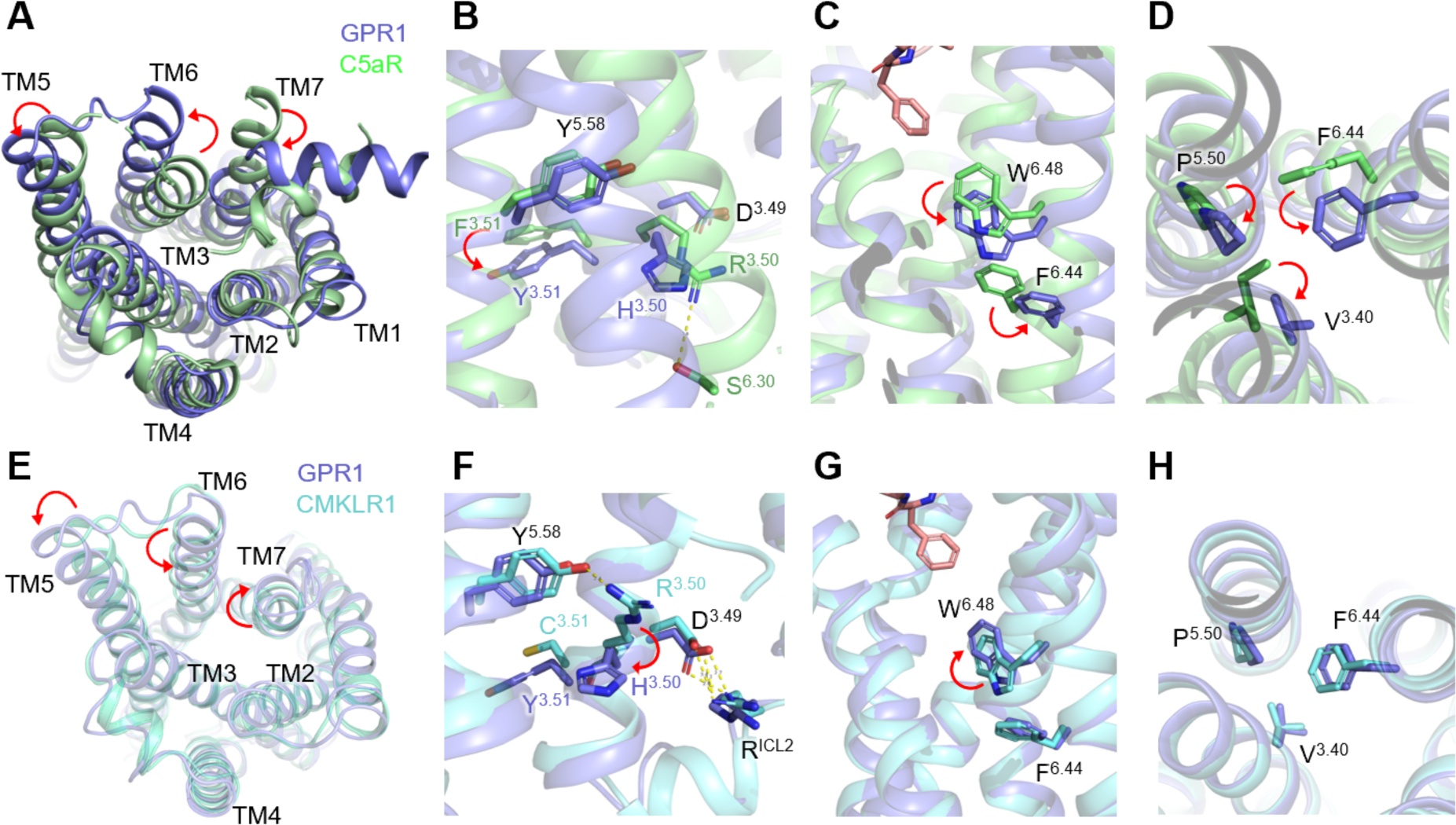
Comparison of GPCR structural motifs for G protein activation. **(A)** Intracellular view of the movement of GPR1 transmembrane helix 5, 6, and 7 (shown in marine blue) in comparison with inactive C5aR (PDB ID: 6C1R, shown in lime green). **(B)** Side close-up view of the D^3 49^-R^3 50^-Y^3 51^ motif. A downward movement of Y^3 51^ of GPR1 is highlighted by a red arrow. (C) Side close-up view of the “toggle switch”, W^6 48^ and F^6 44^, an anti-clockwise rotation is highlighted for GPR1. (D) Rotamer conformational changes at the P^5^ 50-I/V3 ^4^°-f6 ^44^ motif of GPR1 and C5aR, respectively. (E) Intracellular view of the movement of GPR1 transmembrane helix 5, 6, and 7 (shown in marine blue) in comparison with active CMKLR1 (PDB ID: 7YKD, shown in cyan). (F) Side close-up view of the D^3 49^-R3 5o.y3.5i _mo_tif downward movement of H^3 50^ of GPR1 is highlighted by a red arrow. **(G)** Side close-up view of the “toggle switch”, W^6 48^ and F^6 44^, a clockwise rotation is highlighted for GPR1. (H) No significant conformational change at P^550^-IA/^3’40^-F^6 44^ _mo_tjf of GPR1 and CMKLR1, respectively.

The highly conserved residue W259^6.48^ as a “toggle switch” of G protein activation showed an anti-clockwise rotation in the GPR1 structure (Fig. 6C), which is characteristic in other activated GPCRs (Weis & Kobilka, 2018; Zhou et al., 2019). For the P^5.50^-I/V^3.40^-F^6.44^ motif, rotamer conformational changes were displayed in GPR1 in comparison with the inactive C5aR (Fig. 6D). To further identify the structural basis for different signaling of GPR1 and CMKLR1 upon activation, we compared active structures of the two receptors. The C9 peptide ligand was found to have an average of 1.2 Å upward shift away from the “toggle switch” W259^6.48^ in GPR1, and the residues in contact with the ligand were closer to the extracellular loop of GPR1 than CMKLR1 (Fig. S7). An outward movement of TM5, TM7 and an inward movement of TM6 was demonstrated in both receptors (Fig. 6E). As for the DRY motif, CMKLR1 presents a DRC in position and R^3.50^ formed a polar interaction with the Y^5.58^ residue (Fig. 6F). H135^3.50^ in GPR1, however, pointed to the cytosolic direction with no observable polar interaction with adjacent receptor residues. For the “toggle switch”, W259^6.48^ in GPR1 shifted slightly upwards (Fig. 6G). There is not much difference observed in the orientation of the P218^5.50^-V125^3.40^-F255^6.44^ motif (Fig. 6H). For residues lining all these examined motifs, the orientation and geometry of side chains in the chemerin-bound GPR1 and C9-bound GPR1 are the same (Fig. S8). Overall, despite differences in the geometry and important motifs between GPR1 and CMKLR1, GPR1 is able to activate the Gi proteins albeit with a lower amplitude.

Next, the interaction between an activated GPR1 and the Gi class of heterotrimeric G proteins was examined. In this study, we adopted DNGαi1, a dominant negative form of human Gαi1 containing the G203A and A326S mutations for decreased affinity for nucleotide binding and increased stability of heterotrimeric G protein complex (Lee et al., 1992; Posner et al., 1998). In the structure, the α5 helix of Gαi inserts into the intracellular loops of GPR1, forming hydrophobic interactions with F76^2.43^, L151^4.39^, V251^6.40^, Y226^5.58^, T247^6.36^, K310^8.49^ (Fig. 7A). Of note, some polar interactions are found between α5 helix G352 and GPR1 H135^3.50^, α5 helix N347 and GPR1 H138^3.53^, αN helix R32 and GPR1 H146^ICL2^ (Fig. 7A). All residues of GPR1 accommodating Gα binding share the same sidechain geometry, implying the conservation of structure in G protein activation by different ligands (Fig. S8). Additionally, we also observed a hydrogen bond between GPR1 helix 8 and D312 of the Gβ subunit (Fig. 7B).

**Figure 7.**
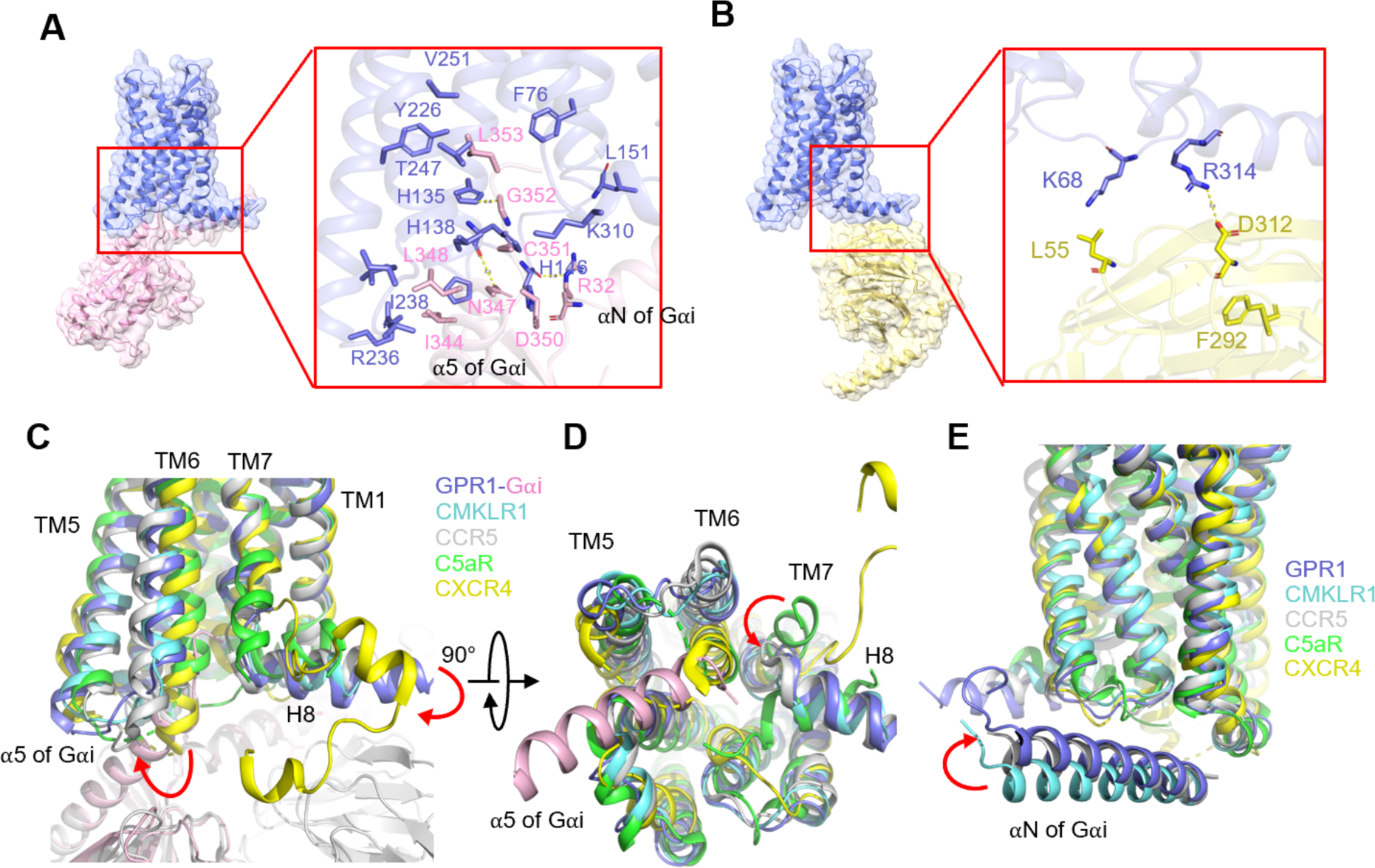
G protein interface of the C9-bound GPR1-Gi complex. **(A)** The interactions between the a5 helix of Gai (pink) and GPR1 (marine blue) in the cavity at ICL3, TM5, TM6, and TM7 regions. (B) The interactions between Gp subunit (yellow) and H8 of the receptor (marine blue). **(C)** Comparisons of the interactions between the a5 helix of Gai and TM5, TM6, and ICL3 of several Gi-coupled receptors including GPR1 (marine blue), CMKLR1 (cyan, PDB ID: 7YKD), CCR5 (gray, PDB ID: 7F1R), C5aR (lime green, PDB ID: 6C1R) and CXCR4 (yellow, PDB ID: 3ODU). **(D)** 90°orientation of (C) for intracellular view showing the locations of ICL2, ICL1, and H8. (E) Same as (C) and (D) yet the interactions of the aN helix of Gai with these receptors are compared.

**Figure 8.**
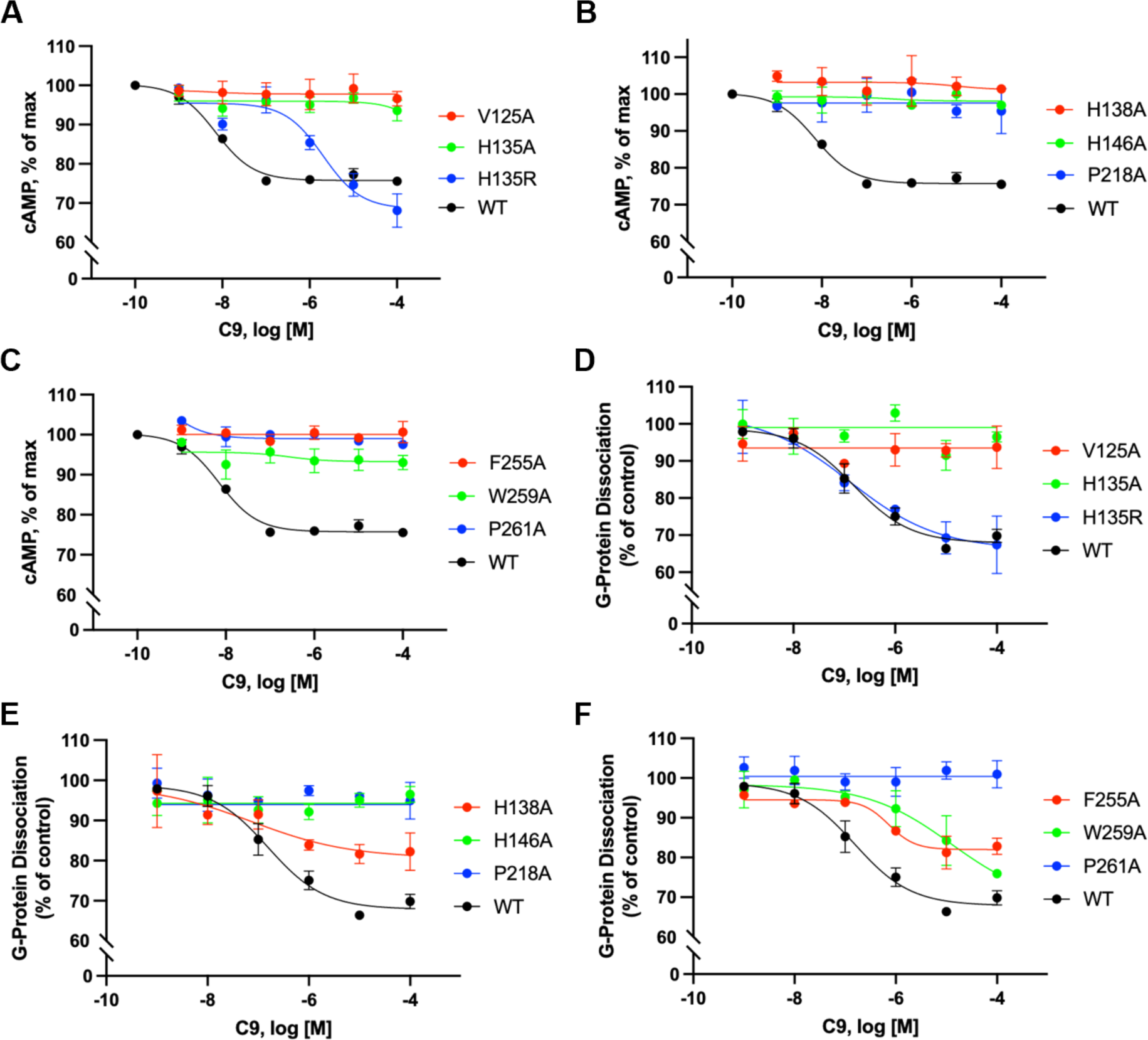
Point mutations at key residues for G protein activation affect cAMP inhibition and G protein dissociation. **(A - C)** cAMP response in HeLa cells transfected to express WT or mutant GPR1. Different concentrations of C9 are applied. **(D - F)** G protein dissociation in HEK293T cells co-transfected to express WT or mutant GPR1, Gai1-LgBiT, GP1, and SmBiT-Gy2. Different concentrations of C9 are applied. All data shown are means ± SEM from three independent experiments.

The interaction between the heterotrimeric Gi protein and the receptor was next compared with several Gi-coupled GPCRs, including the active CMKLR1 and CCR5 (PDB ID: 7F1R), and the inactive C5aR and CXCR4 (PDB ID: 3ODU) (Fig. 7 C-E). The orientation of TM6 and TM7 marks the most remarkable difference between active and inactive receptors (Fig. 7C and D). For the active receptors GPR1, CMKLR1 and CCR5, TM6 displays an outward tilt allowing space for an interface between the receptor and the C-terminal α5 helix of Gαi (Fig. 7C). Helix 8 of the active receptors shows a movement to the intracellular compartment for engagement of the Gβ subunit (Fig. 7C). An inward movement of TM7 is also observed for GPR1 and other active receptors compared with the inactive representatives (Fig. 7D). Although we did not observe much polar interaction between GPR1 and the αN helix of Gαi as in CMKLR1 (Wang et al., 2023), the αN helix of Gαi has moved upwards for closer proximity with the receptor helix 8 (Fig. 7E). These features contribute to the activation of G protein by GPR1.

We verified the proposed mechanisms of G protein activation by introducing point mutations to the key residues. In addition to the substitution of H135^3.50^ into the canonical arginine, as mentioned above and shown in Fig. 7A, point mutations of P218^5.50^-V125^3.40^-F255^6.44^ into alanine greatly reduced the cAMP inhibition as well as G protein dissociation (Fig. 8). Substitution of the “toggle switch” W259^6.48^ with alanine caused a complete loss in cAMP inhibition, yet G protein dissociation was partially retained with a decreased efficacy and potency (Fig. 8C and F). For the interaction interface between the receptor and Gαi, alanine substitution at H135^3.50^, H138^3.53^ and H146^ICL2^ completely diminished G protein dissociation and cAMP inhibition (Fig. 8 A-B and D-E). All these functional data confirmed the importance of the aforementioned key residues in activating Gi proteins.

## Discussion

There is increasing awareness of the importance of chemerin in immune regulation and lipid biogenesis, that are biological processes associated with a number of common illnesses including acute inflammation, psoriasis, angiogenesis, obesity, diabetes, nonalcoholic fatty liver disease, ovarian cancer and endometrial cancer (Bozaoglu et al., 2007; Ernst & Sinal, 2010; Goralski et al., 2019; Goralski et al., 2007; Helfer & Wu, 2018; Kennedy & Davenport, 2018; Macvanin et al., 2022; Parolini et al., 2007; Schioppa et al., 2020; Su et al., 2021; Sun et al., 2021; Yue et al., 2023). It is therefore necessary to understand how chemerin exerts these physiological and pathological functions through the chemerin receptors. Of the three chemerin receptors identified, CMKLR1 is well known for mediating most biological activities of chemerin, and CCRL2 is a non-functional chemerin receptor serving as a scavenger (Kennedy & Davenport, 2018). In comparison, the functions of GPR1 are less clear. Initially identified as an orphan receptor (Marchese et al., 1994), the natural ligand of GPR1 was not clear until 2008, when Barnea *et al* reported chemerin-induced membrane association of β-arrestin2 using a functional assay for protein-protein interaction (Barnea et al., 2008). Chemerin was more potent on GPR1 (EC_50_ of 240 pM) than on CMKLR1 (3 nM) in β-arrestin2 recruitment in this assay. The same also applied to the C9 peptide, that showed a higher potency (EC_50_ of 1 nM) on GPR1 than on CMKLR1 (24 nM). Based on these results, GPR1 was considered a receptor that preferentially activates the β-arrestin2 pathway (Barnea et al., 2008; Fischer et al., 2021), although other studies showed its ability to mediated G protein signaling as well (De Henau et al., 2016; Degroot et al., 2022).

In this study, we first established that GPR1 functionally couples to the Gi proteins with very low potency in β-arrestin signaling. These results led us to prepare the chemerin-GPR1-Gi complex (and C9-GPR1-Gi complex) for structural studies using cryo-EM. Our findings represent the first high-resolution structure of full-length chemerin bound to one of its receptors, that allowed us to make comparison of this chemotactic small protein with chemokines and chemotactic peptides. Our findings also confirmed an early prediction that chemerin acts as a “reverse chemokine” with an N-terminal core region and a C-terminal loop for receptor activation, which is the mirror-image of a canonical chemokine that uses its N-terminus for receptor activation and C-terminus as core regions (Zabel et al., 2006). This mode of binding, demonstrated experimentally in our study, may explain why chemerin bind to the C-C chemokine receptor CCR2 without eliciting any downstream signaling activity (Zabel et al., 2008). Another piece of evidence that supports the role of chemerin as a chemokine-like chemoattractant (Kumar et al., 2019; Zabel et al., 2006) is that chemerin uses a “two-site” model that is similar to chemokine interaction with chemokine receptors (Kennedy & Davenport, 2018; Siciliano et al., 1994). Notably, our cryo-EM structure of chemerin-bound GPR1 Gi protein signaling complex provides direct proof for such a “two-site” model for chemerin binding with the following features. At chemerin binding region 1 (CBR1), multiple polar and non-polar interactions between the N-terminus of GPR1 (CBR1) and the cavity formed between H1 helix and β1 sheet of the N-terminal core of chemerin stabilize the chemerin binding. The importance of GPR1 N-terminal CBR1 in stable chemerin binding is reported recently using chimeric receptor-based binding assays (Kretschmer et al., 2023). At chemerin binding region 2 at the receptor TM pocket, contacts between the C-terminus of chemerin (C-terminal nonapeptides, structurally identical to C9 peptide) and CBR2 residues activate GPR1 downstream signaling. In some chemokine receptors, the receptor ECL2 form polar contacts with chemokine ligands, which defines a chemokine recognition site 3 (CRS3) (Urvas & Kellenberger, 2023). Similar feature presents in chemerin-GPR1 interactions, where E129 at H2 helix of chemerin forms hydrogen bond with H194^5.26^ of GPR1 close to ECL2 region. This region is therefore defined as CBR3 for chemerin binding. Canonical chemokine recognition sites involve a structurally-conserved site 1.5 (CRS1.5), where the N-terminus of the receptor interacts with the CC/CXC motif of the chemokine (Kleist et al., 2016; Urvas & Kellenberger, 2023). This interaction, however, is absent in chemerin-GPR1 structure. It is concluded that chemerin, which is not a chemokine, has retained some of the features that characterize the chemokine-chemokine receptor interaction.

Our structural model also provides molecular insights into the mode of binding between the C-terminal nonapeptide of chemerin (C9) and GPR1, specifically at CBR2. C9 takes an “S”-shape pose in the binding pocket of GPR1, which is surrounded by TM2, 3, 4, 6, 7 and deep enough to accommodate the nonapeptide. The C terminus of S9 reaches the bottom of the binding pocket and forms extensive polar interactions with S114^3.29^, Q118^3.33^ and R176^4.64^. This model differs from the proposed model by Fischer *et al* that was developed through homology modeling and molecular docking (Fischer et al., 2021). In the Fischer model, the C9-GPR1 interaction is dominated by hydrophobic interactions between F8 of the ligand and a hydrophobic domain in the extracellular loop 2 of GPR1 consisting of F^4.69^, L^4.74^, Y^4.76^, and F^4.79^. This feature, combined with a proposed hydrophobic interaction between F6 and F^2.68^, renders a shallow hydrophobic pocket for ligand binding involving mostly the extracellular domains as seen in many chemokine receptors. For ligand accommodation in the Fischer model, the peptide ligand bent between G4 and Q5, pointing towards T^7.39^. Apparently, the Fischer model is not energetically favorable for stable binding of the C terminal peptide, which is critical to activation of the receptor for G protein signaling.

Our structural model provides direct evidence for an interaction of GPR1 and the Gi proteins, that clarified an issue on whether GPR1 is able to signal through G proteins. Analysis of the receptor-Gi protein interface found some polar interactions between the receptor and G protein heterodimer complex. Of note, the highly G protein-binding motif, D^3.49^-R^3.50^-Y^3.51^, found in many Class A GPCRs, is replaced by DHY in GPR1. Surprisingly, histamine substitution in this motif did not diminish G protein binding. Notably, a polar interaction between H135^3.50^ and α5 helix of Gi is evident. This structural feature indicates that the canonical DRY motif is not absolutely necessary for GPCR-Gi interaction, and the actual coupling of the receptor to G protein may be context-specific in each of the receptors.

The chemerin/C9-bound GPR1-Gi signaling complex structure was compared with the C9-CMKLR1-structure that we resolved earlier (Wang et al., 2023). There are high degrees of homology at both primary sequence level (71% sequence homology, with 37% identical amino acids) (Kennedy & Davenport, 2018) and 3D structure level. For G protein interacting interface, CMKLR1 has additional polar interactions with the αN helix of Gαi than GPR1. The difference in DRY motif (DRC in CMKLR1, DHY in GPR1) does not seem to affect Gi coupling in pharmacological term. Another structural feature, the absence of the polar interaction in GPR1 between H^3.50^ and Y^5.58^ (due to the R^3.50^ to H^3.50^ switch), does not produce a functional impact in Gi coupling. Of importance, the overall receptor structure does not change despite the R^3.50^ to H^3.50^ switch. Hence, the lower G protein coupling efficiency in GPR1 may be explained by fewer interactions at the receptor-Gαi interface, but not the R^3.50^ to H^3.50^ switch as predicted before. For the receptor-C9 interaction, the C9 peptide resumes an “S” shape in both GPR1 and CMKLR1. However, C9 insertion into the GPR1 binding pocket is not as deep as it is in CMKLR1, showing an average of 1.2Å upward shift in GPR1. As a result, the bottom of the GPR1 binding pocket is higher above W^6.48^, the ‘toggle switch’ for G protein activation, when compared with CMKLR1 (Wang et al., 2023). Taken together, structural analysis of the GPR1-Gi signaling complex provides evidence that GPR1 is fully capable of activating Gi proteins despite minor sequence differences at the DRY motif and a shallower binding pocket for the peptide ligand.

While the structural information of GPR1 expands our knowledge in chemerin receptor biology, several unknowns still remain for further investigation. Given the high similarities between GPR1 and CMKLR1, they may share a variety of agonists and antagonists. Currently the only natural ligand of GPR1 and CMKLR1 is chemerin, and the wide distribution of GPR1 and CMKLR1 among immune cells, adipose tissues and central nervous system suggests the presence of other possible endogenous ligands (Herova et al., 2015; Marchese et al., 1994; Tokizawa et al., 2000; Wittamer et al., 2003; Wittamer et al., 2004). Indeed, a previous report unraveled a novel ligand of GPR1, FAM19A1, which is highly expressed in adult hippocampus and has neural modulatory effect (Zheng et al., 2018). However, the reported study was based primarily on animal experiments without a biochemical mechanism for receptor activation, and there is still a lack of structural information about FAM19A1 interaction with GPR1. Another unresolved issue is whether GPR1 and CMKLR1 differ in activating the β-arrestin pathway, as currently there is a lack of structure for a ligand-bound receptor-β-arrestin complex. Further investigation in these directions will help to understand how the signals generated by chemerin binding are mediated by these two receptors.

## Acknowledgments

This work was supported in part by grants from the Science, Technology and Innovation Commission of Shenzhen Municipality GXWD20201231105722002-20200831175432002 (R.D.Y., Y.D. and L.Z.), JCYJ20200109150019113 (Y.D., G.C.), JCYJ20200109150003938 (L.Z.), RCYX20200714114645019 (L.Z.) and RCBS20221008093330067 (A.L.). This work was also supported by National Natural Science Foundation of China 31971179 (L.Z.), 32271263 (Y.D.) and 32070950 (R.D.Y.), China Postdoctoral Science Foundation 2022M713049 (A.L.) and 2021M703092 (J. W.), the Ganghong Young Scholar Development Fund (R.D.Y.), the Kobilka Institute of Innovative Drug Discovery at The Chinese University of Hong Kong, Shenzhen (Y.D., R.D.Y.), and the fund from Shenzhen-Hong Kong Cooperation Zone for Technology and Innovation (HZQB-KCZYB-2020056). The authors thank the Kobilka Cryo-Electron Microscopy Center in the Chinese University of Hong Kong, Shenzhen for cryo-electron microscopy analysis, and Warshel Institute for Computational Biology (funding from Shenzhen City and Longgang District) for computational work.

## Author Contributions

R.D.Y., A.L. and Y.L. designed the research; A.L. performed cloning, protein expression and purification, screening of cryo-EM grids, collection of cryo-EM data and building of the structural model with help from G.C. and F.Y.; Y.L. conducted mutagenesis analysis, functional assays, structural comparison; W.L. and L.Z. performed MD simulation; J.W. designed and performed the Nano-BiT assay; Q.L. performed initial modeling and functional assays of C9 peptide; L.Z., Y.D. and R.D.Y. supervised the research; All authors discussed and analyzed the data. Y.L., A.L. and R.D.Y. prepared the figures and wrote the manuscript.

## Competing Interest Statement

The authors declare no competing interest.

---

**Fig. S1.**
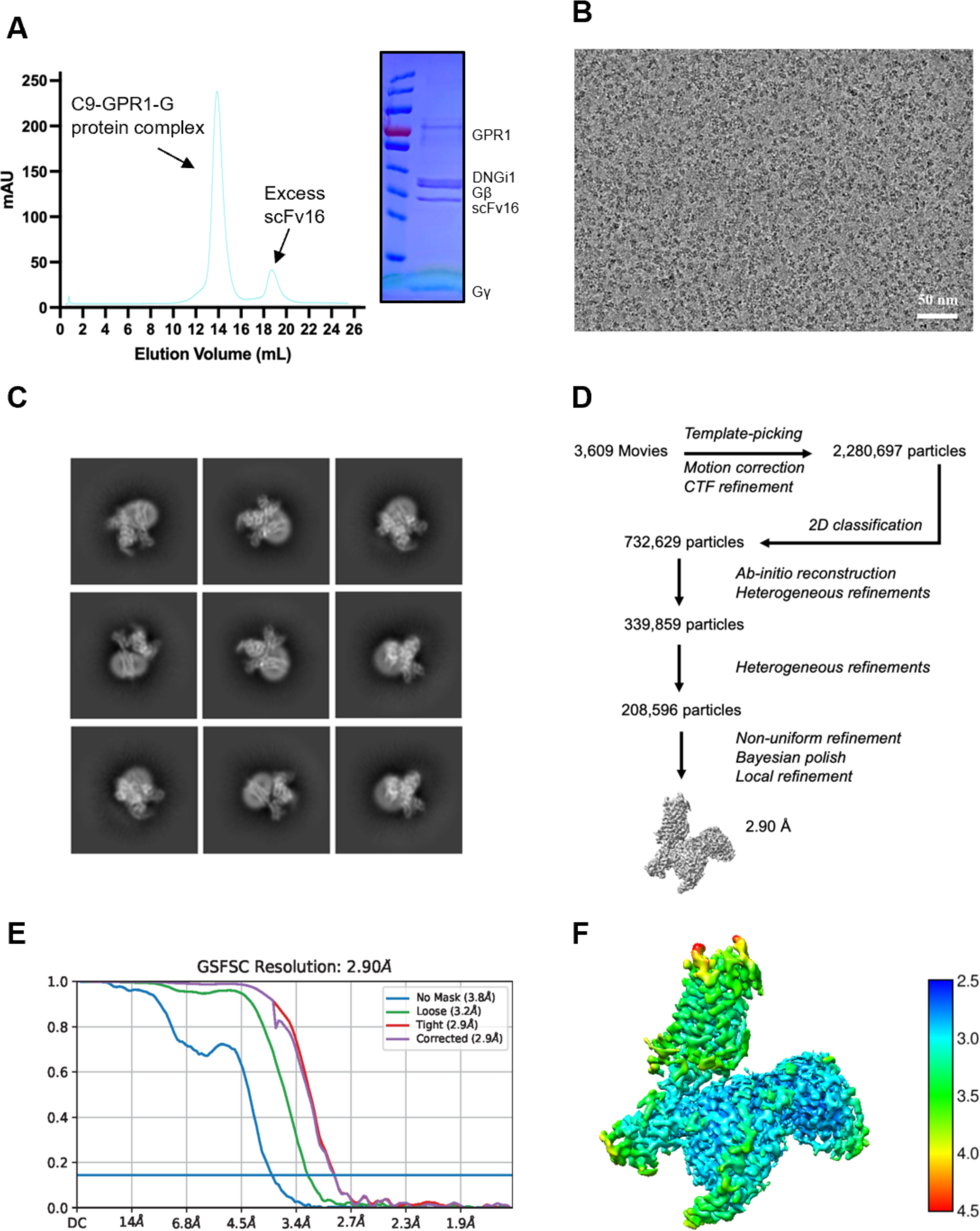
Protein purification, cryo-EM data collection, structure determination and cryo-EM maps. **(A)** Size-exclusion chromatography elution profiles of the C9-GPR1-G protein complex and the SDS-PAGE and Coomassie blue staining of the C9-GPR1-G protein complex. **(B)** Representative micrograph of the complex particles from 3,609 movies. (C) Representative 2D averages. **(D)** Workflow for cryo-EM image processing. **(E)** Gold standard Fourier shell correlation (FSC) curve indicates overall nominal resolution at 2.90 A using the FSC=0.143 criterion. **(F)** Local resolution map.

**Fig. S2.**
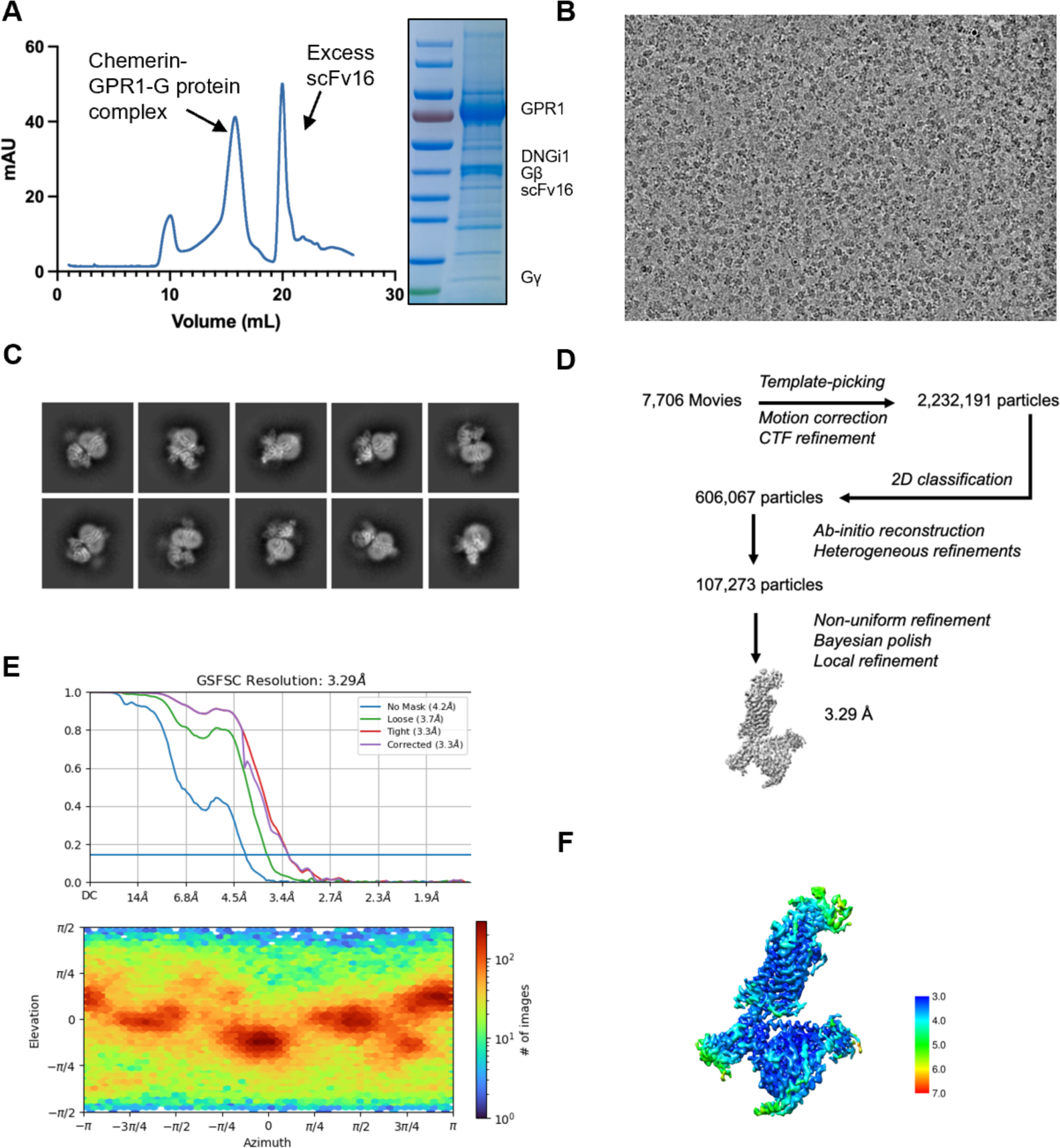
Protein purification, cryo-EM data collection, structure determination and cryo-EM maps. **(A)** Size-exclusion chromatography elution profiles of the chemerin-GPR1-G protein complex and the SDS-PAGE and Coomassie blue staining of the chemerin-GPR1-G protein complex. (B) Representative micrograph of the complex particles from 7,706 movies. (C) Representative 2D averages. **(D)** Workflow for cryo-EM image processing. **(E)** Gold standard Fourier shell correlation (FSC) curve indicates overall nominal resolution at 3.14 A using the FSC=0.143 criterion. (F) Local resolution map.

**Fig. S3.**
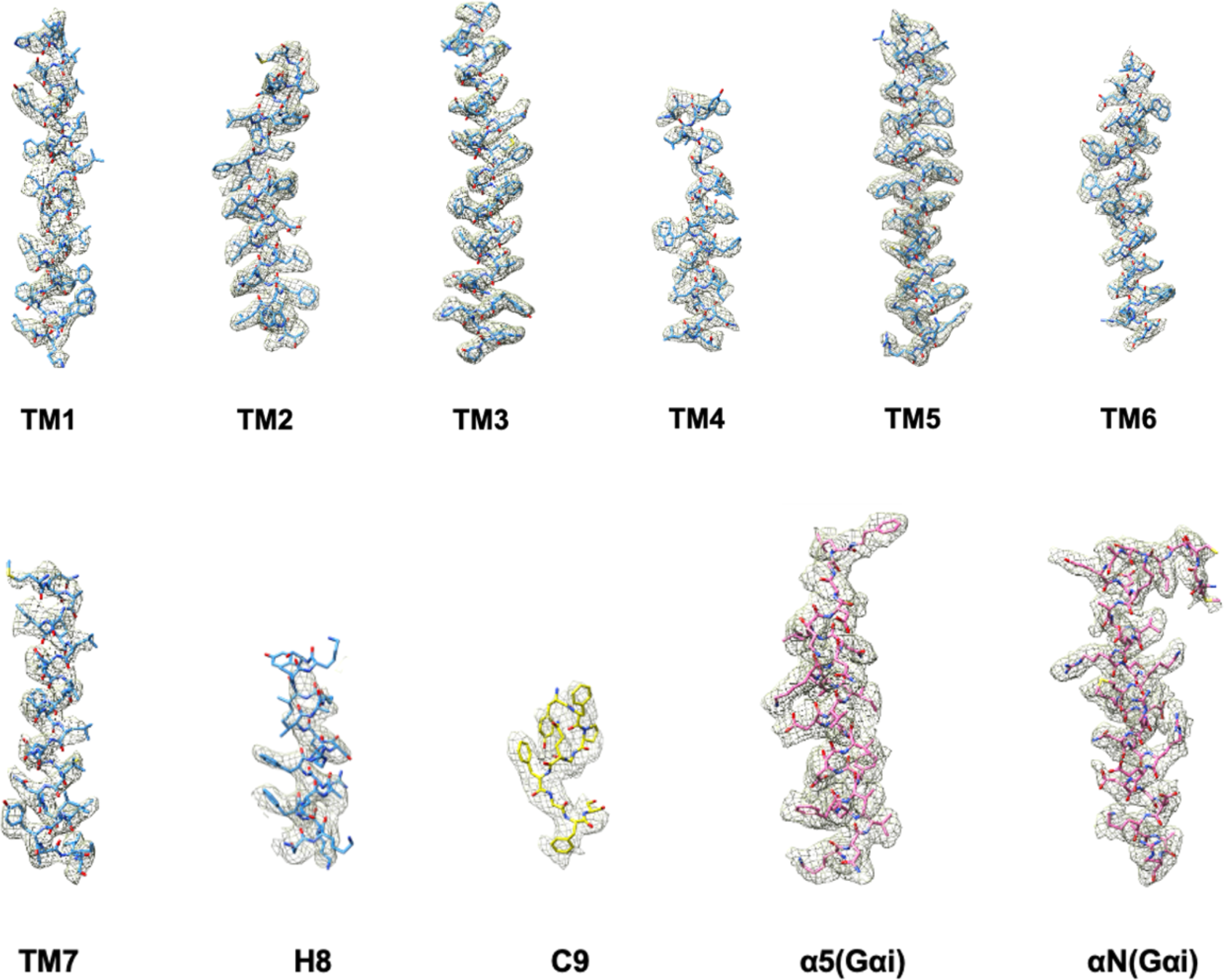
Representative density maps and models for TM1-7 and H8 C-terminal a helices of Gail (a5 and aN) and the ligand C9 peptide.

**Figure S4.**
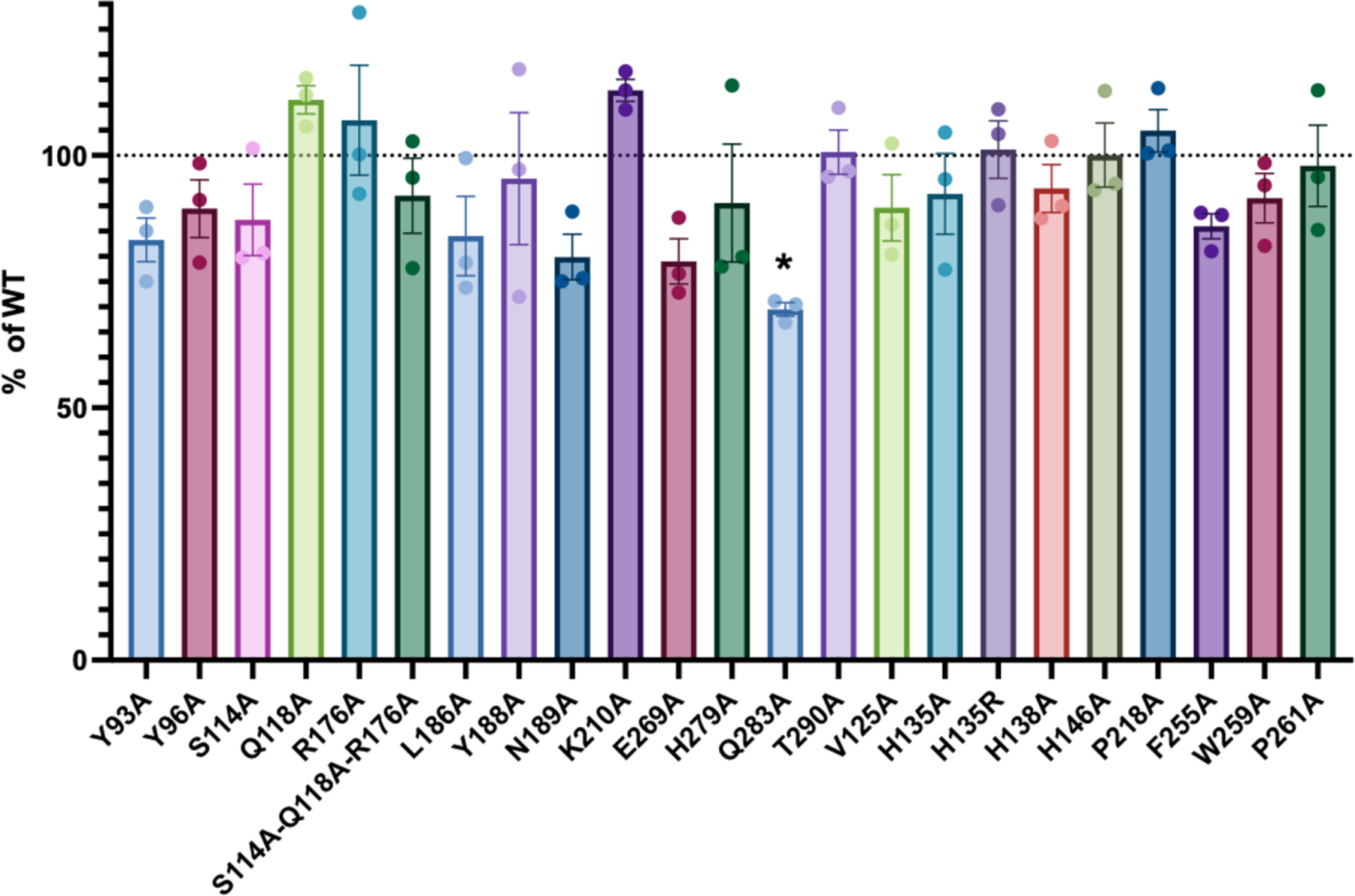
Cell surface expression of mutants. HEK293T cells were transfected with FLAG-tagged WT or mutant GPR1 for 24 h at 37°C. The cells were then incubated with an FITC-conjugated FLAG antibody for 30 min on ice. The fluorescence signals on the cell surface were quantified by flow cytometry. Data shown are means ± SEM from 3 independent experiments. *\*, p <* 0.05.

**Figure S5.**
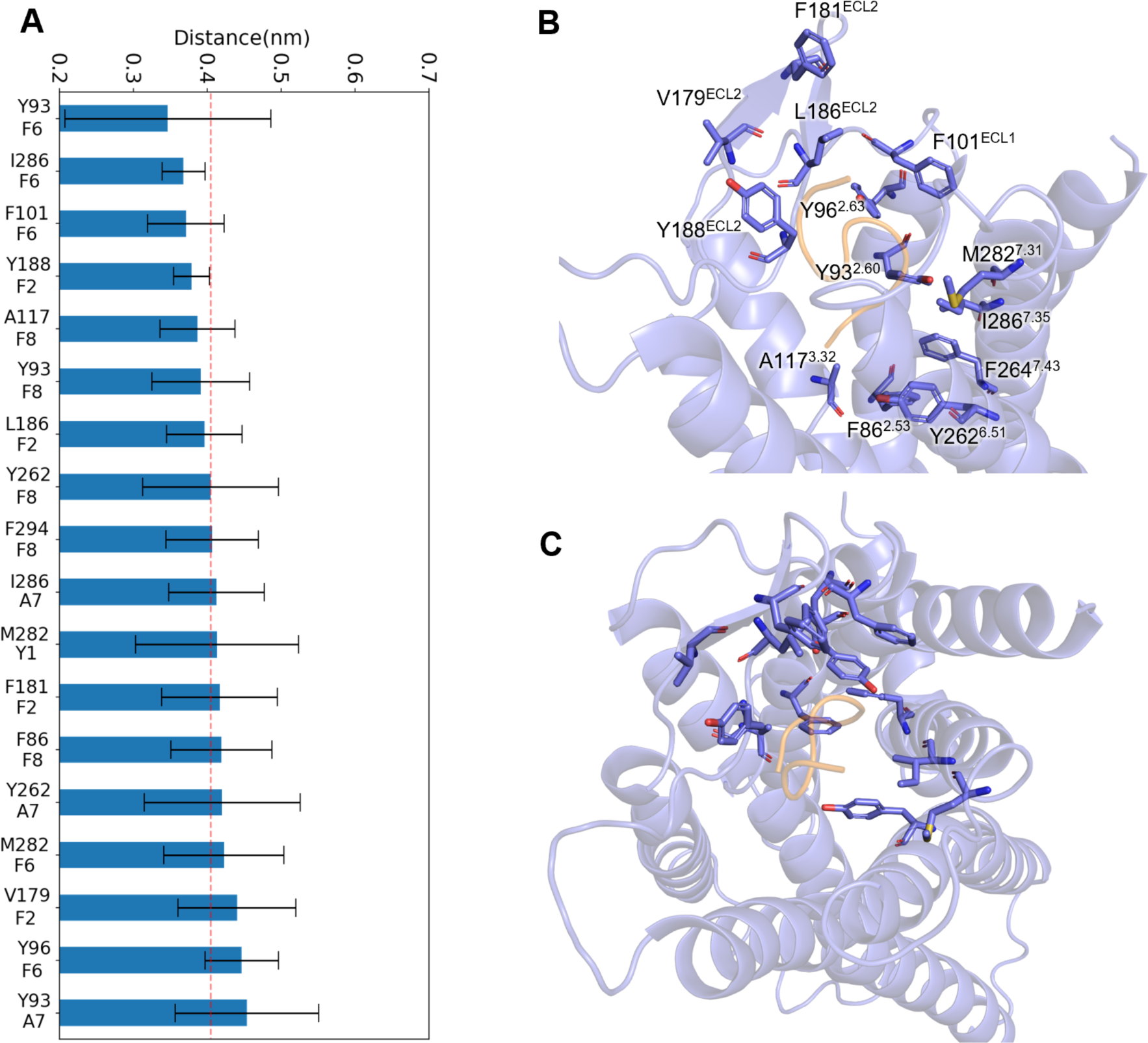
The hydrophobic contact analysis of the C9 peptide binding on GPR1. **(A)** The average heavy-atom distance of close-contact residues between C9 and GPR1. (B-C) The relative distribution of the close-contact residues around the C9 peptide.

**Figure S6.**
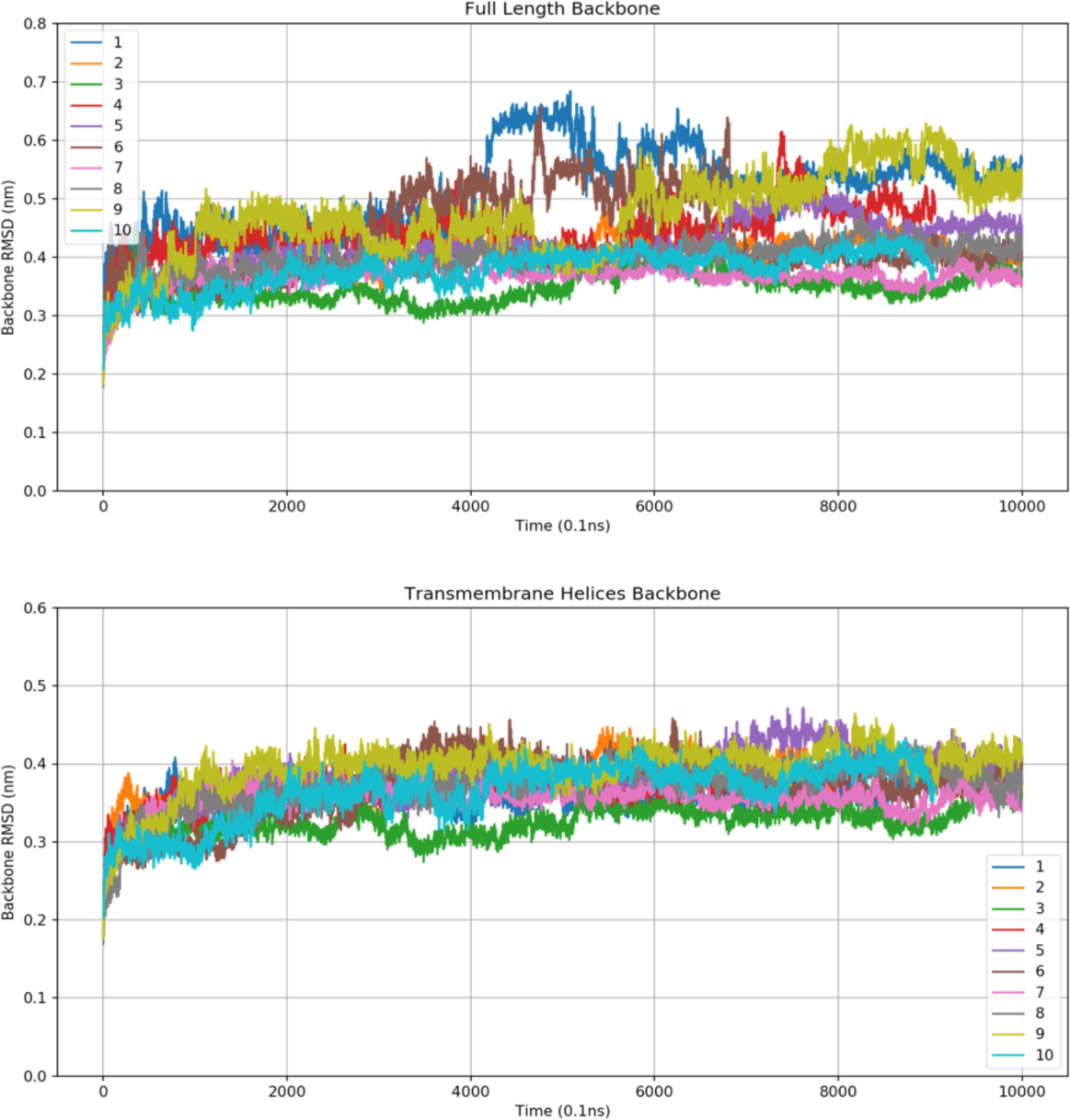
Backbone RMSD of the full length (upper panel) and trans­membrane helices (lower panel) of GPR1 in each replica of MD simulation.

**Figure S7.**
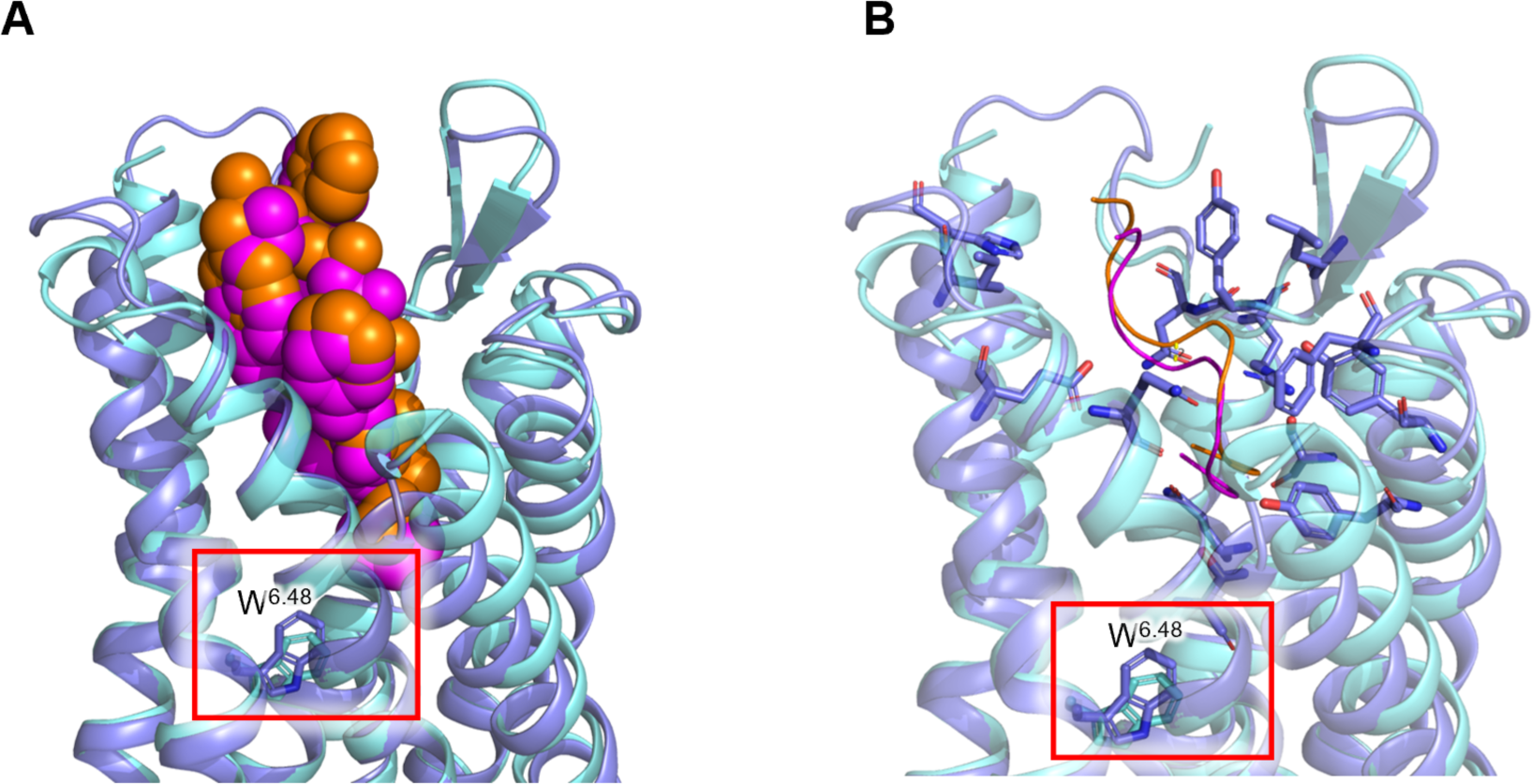
Comparison of the ligand binding pocket of GPR1 (marine blue) and CMKLR1 (cyan). **(A)** Sphere representation of C9 bound to GPR1 is shown in orange, and the C9 bound to CMKLR1 is shown in magenta. W^648^ “toggle switch” is highlighted with stick representation. (B) Key interaction residues of the binding pockets of GPR1 and CMKLR1 are shown in licorice representation. The C9 ligands respectively bound to the two receptors are shown as ribbon. The distance of the ligands is calculated and shown as a centroid distance of 1.2 A.

**Figure S8.**
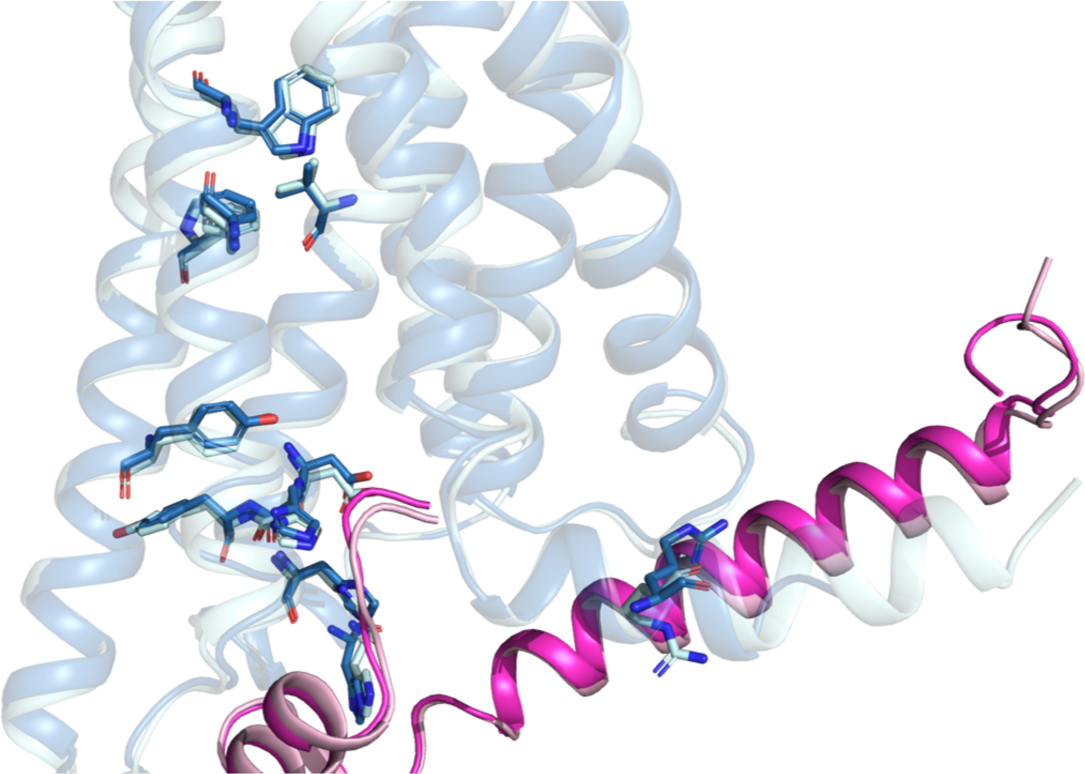
Comparison of structural motifs for G protein activation between GPR1 (dark blue) and CMKLR1 (cyan). The residues responsible for G protein activation are highlighted in sticks. The G protein in the GPR1-Gi structure is shown in magenta, and the G protein in the CMKLR1-Gi structure is shown in light pink.

**Table S1.**
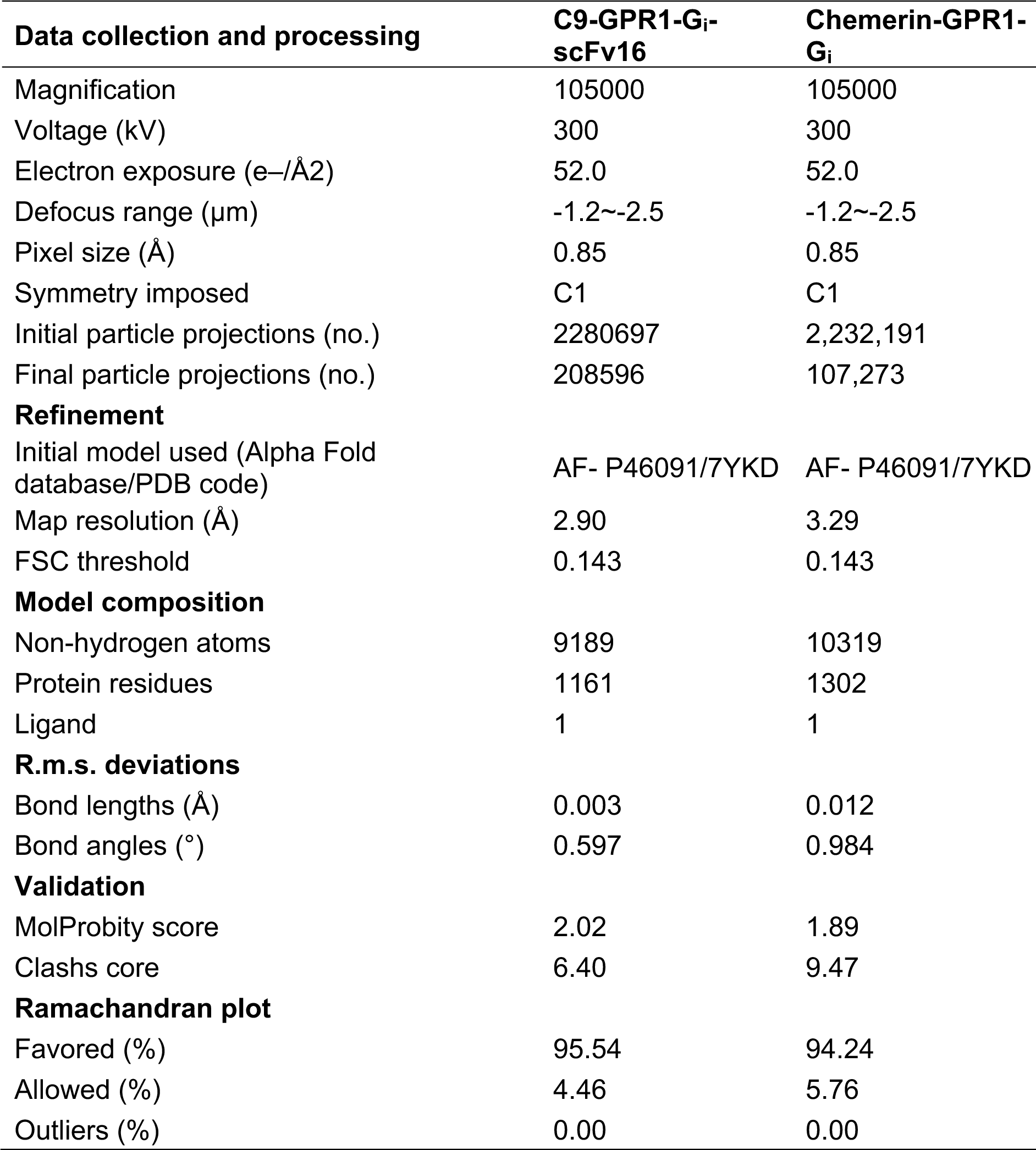
Cryo-EM data collection, model refinement and validation statistics.

| **Data collection and processing** | **C9-GPR1-G<sub>i</sub>-scFv16** | **Chemerin-GPR1-G<sub>i</sub>** |
| --- | --- | --- |
| Magnification | 105000 | 105000 |
| Voltage (kV) | 300 | 300 |
| Electron exposure (e-/Å<sup>2</sup>) | 52.0 | 52.0 |
| Defocus range (μm) | -1.2~-2.5 | -1.2~-2.5 |
| Pixel size (Å) | 0.85 | 0.85 |
| Symmetry imposed | C1 | C1 |
| Initial particle projections (no.) | 2280697 | 2,232,191 |
| Final particle projections (no.) | 208596 | 107,273 |
| **Refinement** |  |  |
| Initial model used (Alpha Fold database/PDB code) | AF- P46091/7YKD | AF- P46091/7YKD |
| Map resolution (Å) | 2.90 | 3.29 |
| FSC threshold | 0.143 | 0.143 |
| **Model composition** |  |  |
| Non-hydrogen atoms | 9189 | 10319 |
| Protein residues | 1161 | 1302 |
| Ligand | 1 | 1 |
| **R.m.s. deviations** |  |  |
| Bond lengths (Å) | 0.003 | 0.012 |
| Bond angles (°) | 0.597 | 0.984 |
| **Validation** |  |  |
| MolProbity score | 2.02 | 1.89 |
| Clashes core | 6.40 | 9.47 |
| **Ramachandran plot** |  |  |
| Favored (%) | 95.54 | 94.24 |
| Allowed (%) | 4.46 | 5.76 |
| Outliers (%) | 0.00 | 0.00 |

## Notes

### Competing Interest Statement

The authors have declared no competing interest.

### Summary of Updates

An additional structure of GPR1-Gi complex bound to chemerin has been included in this updated version of the manuscript. Analysis of both C9-bound and chemerin-bound GPR1 structures has been updated.

